# Comparisons of two receptor pathways in a single cell-type reveal features of signalling specificity

**DOI:** 10.1101/2023.07.03.547518

**Authors:** Yan Ma, Isabelle Flückiger, Jade Nicolet, Jia Pang, Joe Dickinson, Damien De Bellis, Aurélia Emonet, Satoshi Fujita, Niko Geldner

**Affiliations:** Department of Plant Molecular Biology, Biophore, UNIL-Sorge, University of Lausanne, 1015 Lausanne, Switzerland; Department of Fundamental Microbiology, Biophore, UNIL-Sorge, University of Lausanne, 1015 Lausanne, Switzerland; Max Planck Institute for Plant Breeding Research, Cologne, North Rhine-Westphalia, 50829, Germany; Plant Science Research Laboratory (LRSV), UMR5546 CNRS/Université Toulouse 3, 24 chemin de Borde Rouge, 31320 Auzeville Tolosane, France

**Keywords:** Signalling specificity, MAPK/MPK cascade, single cell-type, FLS2, SGN3/GSO1, RLK pathways, MYB36, endodermis

## Abstract

To respond appropriately to diverse stimuli, cells harbour numerous receptor pathways yet share downstream signalling components. Mitogen-Activated Protein Kinase (MPK) cascades, present in all eukaryotes, act as central hubs shared amongst diverse signalling pathways. How signalling specificity is maintained within a single plant cell-type is not well understood. We engineered a genetic background for direct comparisons of a developmental and an immunity pathway in the Arabidopsis root endodermis. We show the two pathways maintain distinct outputs despite similar MPK phosphorylation patterns. Systematic activation of all individual MPK Kinases (MKKs) in the endodermis revealed that different MKK groups can drive distinct outputs. We propose that specificity is achieved through a balance of combinatorial activation and cross-pathway inhibition within the MPK cascade. Our findings demonstrate the ability of MPK cascades to generate diverse outputs in one plant cell-type, providing new insights into the mechanisms contributing to signalling specificity.

## Introduction

Multicellular organisms rely on individual cells to detect and integrate exogenous and endogenous stimuli and translate them into various biological responses. Diverse signalling pathways must accurately maintain the correspondence between a given extracellular input with its appropriate cellular response output. Decades of research have revealed the chain of molecular events from extracellular ligand perception by cell surface receptors to transcriptional regulation in the nucleus. However, a key question remains: how is pathway-specific information maintained all the way to the nucleus when multiple receptor pathways are stimulated within a crowded cell? This problem is sharpened when intracellular events activated by different pathways frequently converge on a limited number of common signalling intermediates. One prominent example is the Mitogen-Activated Protein Kinase (MPK) cascade, a key intermediate module that is activated by numerous stimuli in plants, animals, and fungi.^1–4^ Understanding the mechanisms underlying specificity in such an hourglass architecture, where diverse inputs and outputs converge, presents a significant challenge.

Plants face an amplified “hourglass problem” as they possess a higher number of receptors per cell-type compared to their metazoan counterparts, but nevertheless share extensive downstream signalling components. Arabidopsis, for instance, displays over 200 Leucine-rich-repeat receptor-like kinases (LRR-RLKs),^5, 6^ dwarfing the 11 Toll-like receptors found in humans,^7^ both representing a subset of their total receptor complement. Moreover, despite difficulties in strictly defining cell-types, there is little doubt that humans have many more cell-types than Arabidopsis,^8, 9^ further enhancing the potential complexity of receptor pathways in plants on a per cell basis. In plants, the prevailing notion suggests that common pathway components mediate specific outputs by activating different sets of transcription factors available in different cell-types. Yet, such explanation implies that a single cell-type would accommodate only one receptor/ligand pathway, which is implausible for plant cells. Therefore, investigating how different receptors within a single plant cell maintain signalling specificity becomes even more crucial, compared to highly specialized animal cells. However, such studies are scarce due to the challenges of isolating and analysing signalling events at single-cell resolution, as well as the lack of precise cellular readouts.

To address these challenges, we leverage the highly ordered and linear differentiation of transparent Arabidopsis roots, which provides a unique opportunity to directly observe single cells within an intact organism. Within this context, we establish the Arabidopsis root endodermis as a cellular model to compare two well-studied LRR-RLK receptor pathways at single-cell level (**Figure 1A**). One pathway is an immunity pathway mediated by the FLAGELLIN SENSING2 (FLS2) receptor, recognizing the microbial pattern flg22.^10, 11^ The other pathway is mediated by the receptor SCHENGEN3 (SGN3, also called GASSHO1 (GSO1)) and endogenous Casparian Strip Integrity Factor 1/2 (CIF1/2) peptides, responsible for Casparian strip (CS) establishment in the endodermis.^12, 13^ The endodermis is a specialized cell layer found in all vascular plants. Each endodermal cell bears a ring-like CS, together forming a highly localized and coordinated membrane-cell wall nexus (**Figure 1D**) analogous to the tight junctions in animal intestinal epithelia.^14^ This CS is composed of hydrophobic lignin impregnating the cellulosic primary cell wall, creating an extracellular diffusion barrier in roots that is vital for selective nutrient uptake.^15, 16^ SGN3 localizes specifically to the endodermal plasma-membrane, whereas the CIF peptides are expressed and processed in the central stele.^12, 17^ The SGN3 pathway acts as a surveillance system for the diffusion barrier, enabling the plant to detect developmental or stress-induced discontinuities in its CS, which manifests as CIF peptide leakage. During early root differentiation, the SGN3 pathway drives CS maturation, characterized by the fusion of its membrane domains (**Figure 1A, D**). In situations of chronic or strong stimulation, such as persistent peptide leakage in barrier-defective mutants or strong external peptide treatment, the SGN3 pathway induces ectopic lignification of cell corners, and excess suberisation, which acts as compensatory repair mechanisms for the endodermal barrier (**Figure 1A**).^18–20^ We use these precise and distinctive subcellular features of the SGN3 pathway as unambiguous and quantifiable signalling readouts.

**Figure 1.**
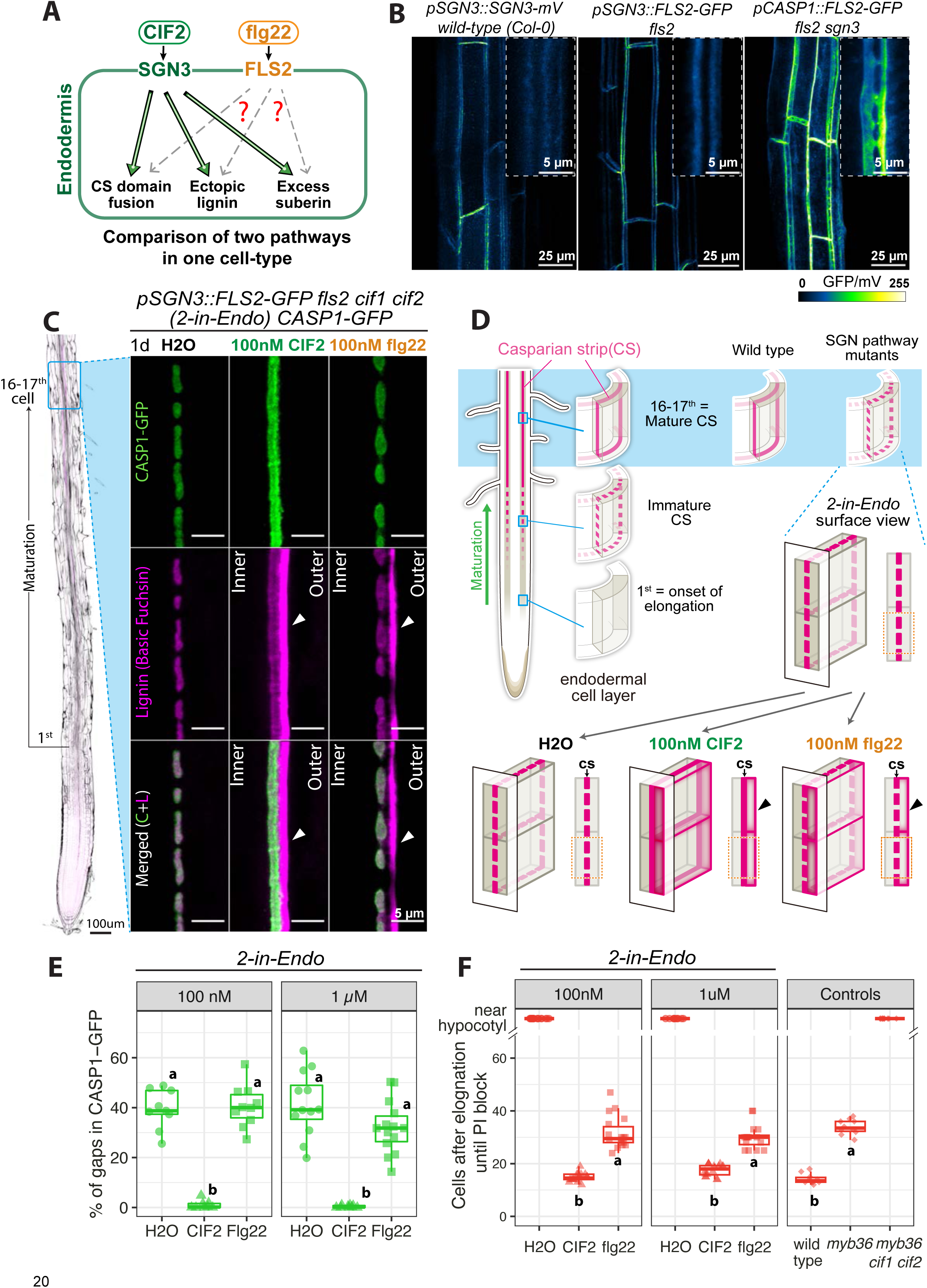
Endodermis-expressed FLS2 cannot functionally replace SGN3 to fuse Casparian Strip (CS) Domains. (A) A single cell-type signalling system in the root endodermis, comparing SGN3/CIF2 developmental pathway and FLS2/flg22 immunity pathway. (B) Endodermis-specific expression and localization of SGN3 (pSGN3::SGN3-mVenus in wild-type), and FLS2-GFP under pSGN3 or pCASP1 in *fls2* or *fls2 sgn3* backgrounds. Representative overviews (maximum projection) and zoomed surface views (see 1D) of mature CS root cells are shown. (C) CS & CASP1 domain fusion in *2-in-Endo CASP1-GFP* after one-day (1d) treatment of H_2_O, 100nM CIF2 or flg22. Untreated *2-in-Endo* root scan (left) outlined by Calcofluor White staining of cellulosic cell walls (black). Endodermal cell surface views (see 1D) at mature CS stage (16-17^th^ cell) show CASP1-GFP (green), lignin stained with Basic Fuchsin (magenta) and overlaps of the two channels (Merged). Arrowheads highlight ectopic lignin. “outer”, cortex-facing, “inner”, stele-facing endodermal surface. (D) Schematics of CS development and 3D architecture. Wild-type root (top-left) shows CS maturation from disconnected domains to fused rings, functioning as a root apoplastic barrier. Early maturation occurs around 11-12^th^ cell (mostly fused), 16-17^th^ cell reaches full maturity (fully fused). SGN3 pathway mutants (right) cannot fuse the CS domains and lack a functional barrier. *2-in-Endo* (bottom) fuses the CS domains in response to CIF2, but not to flg22, yet both trigger ectopic lignin at the cortex side. Illustrations modified from Fujita *et al*.^1^ (E) Quantifications for (C) showing the percentage CASP1-GFP gaps in *2-in-Endo* in response to 1d-treatment with H_2_O, 100nM and 1µM CIF2 or flg22 (n≥8). (F) Propidium Iodide (PI) penetration assay quantifies CS barrier function in *2-in-Endo* after 1d treatments compared to controls (n≥10). Roots with no barrier show PI penetration near root-hypocotyl junction, shown in the “near hypocotyl” category and are excluded from numerical statistical test. For (E)(F), groups with the same letter are not significantly different (P<0.05, one-way ANOVA and Tukey test). Unbalanced Tukey test for unequal replication. See also Figure S1, Movie S1

While the SGN3 and FLS2 pathways are regarded to have markedly different functional outputs (a precise developmental program versus a broad bacterial immune response), there are striking similarities in the signalling structure and components of these pathways. Both receptors recruit co-receptors from the SERK family at the plasma membrane to bind directly and specifically to their respective ligands, flg22 and CIF2, via cognate extracellular LRR domains.^21–23^ These receptor complexes activate members of the Receptor like cytoplasmic kinases VII, namely SGN1 and BIK1 for SGN3 and FLS2 respectively,^20, 24^ which in turn triggers a series of intracellular signalling events. These include downstream signatures common to both pathways, such as Reactive Oxygen Species (ROS) production and activation of MPKs.^20, 25–28^ How are the different outputs achieved when pathways share such extensive similarities? Output differences can be conveniently assigned to the endodermis-specific expression of SGN3 and the broad and dynamically changing expression of FLS2.^29^ However, endodermal cells can also respond to flg22 stimulation,^30^ which prompted us to sharpen the question of signalling specificity in plants by restricting FLS2 expression exclusively to the endodermis and to address the important question: what happens when both pathways are present in a single cell-type?

Here, we present the development of a tailored genetic background that enables the stimulation of the two distinct receptor pathways (FLS2 and SGN3) in the same cell-type — the differentiating root endodermis. Our findings provide clear evidence for the maintenance of signalling specificity in a single cell despite strong pathway activation. We identified the transcription factor MYB36, a crucial regulator of endodermal differentiation,^19, 31^ as a key hub driving pathway-specific outputs in the nucleus. Through extensive, cell-type specific loss and gain-of-function studies, focused on the MPKs and MPK Kinases (MKKs) in the endodermis, we reveal their distinct capacities for transducing both common and specific branches of the two signalling pathways. An important message of our discoveries is the remarkable potential of so-called “common intermediates” such as the elements of the MPK cascades, to produce a wide range of outputs through combinatorial activation of common and specific components. Based on our study, we propose a model in which signalling specificity to the nucleus can be encoded through the combinatorial activation of specific subsets of MAP kinases, highlighting the complex nature of single cell signal decoding in plant cells.

## Results

### Exclusively endodermis-activated FLS2 induces ectopic lignin but cannot functionally replace the endogenous SGN3 pathway to fuse Casparian strip domains

To study signalling specificity at a single cell level, we developed a genetic background that allows stimulation and direct comparison of two distinct pathways in one cell-type — the differentiating root endodermis (**Figure 1A**). In an *fls2* mutant background, we first expressed FLS2 under the endodermis-specific SGN3 promoter (pSGN3::FLS2-GFP) for endodermal FLS2 expression and localisation, mimicking SGN3 (**Figure 1B**). We then knocked out CIF1 and CIF2 genes using CRISPR-Cas9, ensuring that SGN3 signalling only occurs upon external peptide stimulation. We termed the resulting plant line *pSGN3::FLS2-GFP fls2 cif1 cif2* as *2-in-Endo.* CIF2 peptide treatments on *cif1 cif2* mutant are known to efficiently restore the SGN3 pathway function and re-establish a continuous Casparian strip.^20, 23^ flg22 peptide treatments rapidly activate immune signalling from native^30^ and ectopically expressed FLS2 in the root.^30, 32^ Therefore, peptide ligand application (flg22 or CIF2) to the *2-in-Endo* line allows precisely controlled stimulation of the respective pathways (FLS2 or SGN3). Using this line, we investigated whether or not FLS2 could functionally replace SGN3 function.

A hallmark of the SGN3 pathway output is the growth and connection of the Casparian Strip Domains (CSD) from aligned islands (called “string-of-pearls” stage) into a continuous belt around each endodermal cell (**Figure 1D**).^12, 13, 33^ Casparian Strip Domain Protein 1 (CASP1) is specifically targeted to this domain, which marks and predicts lignin deposition at the CS.^25, 34^ We therefore introduced *pCASP1::CASP1-GFP* (CASP1-GFP) into *2-in-Endo*. Note that FLS2-GFP fluorescence is too weak to be detected with the same settings used to detect CASP1-GFP. We confirmed that CIF2 treatment triggers full connection of CASP1 domains and corresponding CS lignin, leaving no gaps at a stage where CS is fully mature in wild-type roots (16-17^th^ endodermal cell after onset of elongation abbreviated 16-17^th^ cell) (**Figure 1C, D, Movie S1**). In contrast, treatments of flg22 cannot improve CS domain fusion, showing comparable percentage of gap areas compared to controls (**Figure 1E)**. We substantiate this finding by visualising CASP1-GFP from initiation to maturation on a greater scale (**Figure S1A**), and by quantifying discontinuity from overviews to incorporate many more endodermal cells at once (**Figure S1B, C**).

Importantly, this strikingly distinct outcome is not due to a lack of FLS2 functionality, as flg22 treatment induces similar ROS production and ectopic lignification at the outer cortex side as CIF2 treatment (**Figure 1C, S1F**). These are additional SGN3 pathway outcomes observed upon strong CIF2 stimulation.^12, 20^ A third outcome with strong CIF2 treatment is the early (nearer the root tip), excess suberin deposition, also mimicked by flg22 treatment in *2-in-Endo* (**Figure 1A, S1D)**. Therefore, the endodermis-specific FLS2 signalling is able to replicate some outcomes of SGN3 but is unable to mediate the crucial aspect of CASP & CS domain growth and fusion. Even with higher FLS2 receptor levels in a *sgn3* background (*pCASP1::FLS2-GFP fls2 sgn3,* **Figure 1B**), FLS2 stimulation cannot induce CASP domain fusion (**Figure S1G, H**). This suggests that CS domain fusion is insensitive to FLS2 signalling, clearly indicating the presence of a main SGN3 signalling branch that FLS2 stimulation cannot induce.

Blocking of Propidium Iodide (PI) penetration provides a sensitive assay to measure CS functionality — it quantifies the developmental stage of an operational extracellular diffusion barrier by scoring PI signal exclusion from the inner side of an endodermal cell (**Figure S1E**).^15, 16^ In wild-type roots, blocking of PI uptake coincides with the early maturation of a continuous CS, assured by a functional SGN3 pathway (**Figure 1F**, ∼14-15^th^ cell in wild-type). A delayed PI block can occur in the absence of a CS through SGN-dependent compensatory lignification at cell corners, as observed in *myb36* compared to *myb36 cif1 cif2’s* absence of PI block (**Figure 1F**). Note that the PI block in *myb36* is achieved purely by compensatory lignin, as the mutant lacks any CS lignin (**Figure 3B)**,^19^ and this blockage is typically delayed by ∼20 cells compared to wild-type plants with a CS (**Figure 1F**).

We demonstrated that the absence of a diffusion barrier in *2-in-Endo* (H_2_O treatment) can be fully rescued by CIF2, restoring an early functional barrier, as in wild-type (**Figure 1F, S1E**). flg22 treatments activate delayed barriers in *2-in-Endo* similar to *myb36*, indicating that *pSGN3::FLS2*-induced ectopic lignin can compensate barrier defects equally well compared to the endogenous SGN3 pathway in *myb36* (**Figure 1F**). In the overexpressing *pCASP1::FLS2* line the barrier function is further improved by flg22 (**Figure S1I**), likely due to a stronger ectopic lignification (**Figure S1G**). Nevertheless, in both cases, barriers are still significantly delayed compared to CIF2-stimulated *2-in-Endo* or wild-type lines.

Altogether, these data demonstrate that despite being able to induce lignification and suberization in the endodermis, FLS2 signalling cannot functionally replace SGN3 for the growth and fusion of CASP1 & CS domains, indicating that signalling specificity between two similar receptor pathways is maintained within the same cell-type.

### Endodermis-expressed FLS2 and SGN3 activate both common and pathway-specific transcriptional responses

To compare transcriptional outputs of FLS2 and SGN3 pathways solely within the endodermis, we conducted comparative RNAseq analysis of *pCASP1::FLS2 fls2* and wild-type seedling roots treated by flg22 and CIF2, respectively at three timepoints (30, 120, 480 min) **(Figure S2A**, data for CIF2 responses from ref^20^). The receptor mutant controls *sgn3* and *fls2,* displayed minimal gene responses to respective peptides (**Figure 2A**). We found a significant overlap between the SGN3 and the FLS2 pathway, with 469 out of 1262 differentially expressed (DE) genes in common (**Figure 2A**). The most enriched Gene Ontology (GO) categories of both pathways show a high degree of overlap with each other, reflecting general biotic & abiotic stress responses (**Figure 2B**). The two shared functional outcomes that we observed, i.e. lignin and suberin deposition (**Figure 1**), are reflected in the transcriptional profiles: Genes involved in lignin and suberin biosynthesis show matching induction patterns after CIF2 or flg22 treatments in respective backgrounds (**Figure 2C, D**). This commonality cannot be simply attributed to shared cell identity (the endodermis), as a cross analysis of highly flg22-responsive genes in whole seedlings carrying native FLS2^35^ reveals similar induction pattern to the SGN3 and FLS2 responses in the endodermis (**Figure S2B**).

**Figure 2.**
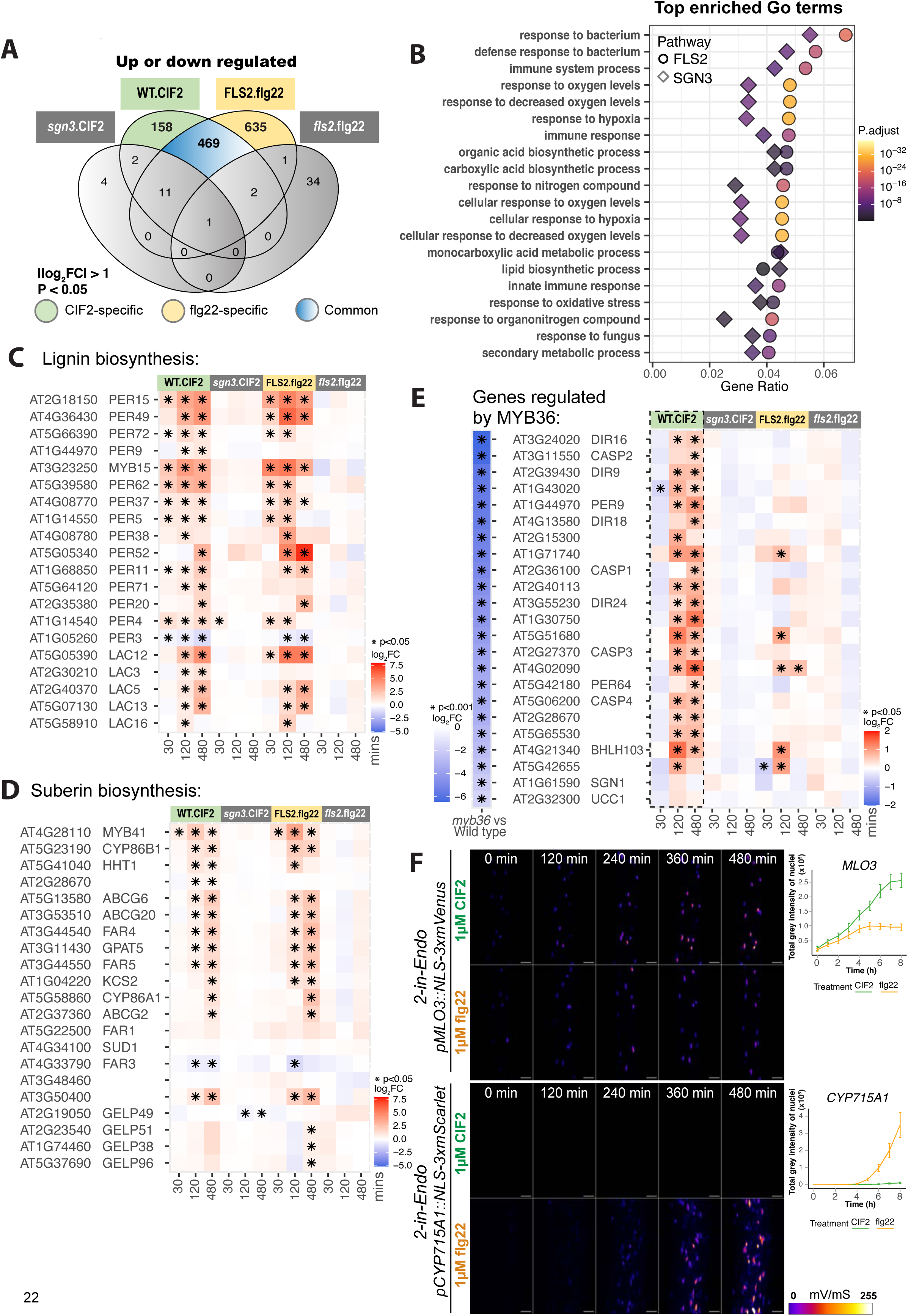
Endodermis-expressed FLS2 and SGN3 trigger common and pathway-specific transcriptional responses. (A) Number of genes significantly regulated by CIF2 or flg22. Cut-off: |Log_2_ fold change (FC)|>1 and P<0.05, for at least one timepoint. Genes that are significantly regulated by one pathway but not the other are considered specific. ‘WT.CIF2’*,‘sgn3*.CIF2’: wild-type/*sgn3* + 1µM CIF2. ‘FLS2.flg22’, *‘fls2*.flg22’: pCASP1::FLS2-GFP *fls2*/*fls2* + 1µM flg22. See Figure S1A for setup. (B) GO enrichment analysis of FLS2 and SGN3 pathway in endodermis. Colour: GO-enrichment significance. **(C-D)** Heatmaps of lignin (**C**) and suberin (**D**) biosynthesis-related gene expression fold changes post-peptide vs H_2_O treatment at indicated time points. (E) Heatmaps of fold changes for MYB36-regulated genes – significantly downregulated in *myb36* compared to wild-type (P<0.001)^2^. For (C-E), asterisks indicate significant regulation (P<0.05). Colour code: degree of fold change. (F) Time lapse images of pMLO3::NLS-3xmVenus (Top) and pCYP715A1::NLS-3xmScarlet (bottom) transcriptional responses in *2-in-Endo* to 1µM CIF2 or flg22. Sum slice projections of roots at 3^rd^-7^th^ cell (top) and 21^st^-25^th^ cell (bottom) shown. Scale bars=25 µm. Quantifications (right panels) show total grey intensity of nuclei signal at each hour (Mean±SE, n=6). See also Figure S2, Table S1, Movie S2, S3.

Despite the overlap, we found 158 and 635 DE genes and GO terms specifically associated with SGN3 and FLS2 pathways respectively (**Figure 2A, S2D**). The greater number of DE genes in the FLS2 pathway could arise from the stronger endodermal CASP1 promoter driving FLS2, compared to the native SGN3 promoter. To validate pathway-specific response genes (**Figure S2C**), we generated transcriptional fluorescent reporters in *2-in-Endo* and found that the majority of reporter lines match the RNAseq predictions (**Figure S2C, Table S1**). For example, *MILDEW RESISTANCE LOCUS O 3* (*MLO3*) shows a much stronger transcriptional response to CIF2 than flg22 (**Figure 2F**). Other SGN-preferential reporter genes include *PRK1*, *LAC3*, *MCA2* (**Table S1, Movie S2**). In contrast, Cytochrome P450 family gene (*CYP715A1*) is a FLS2-specific reporter that responds strongly to flg22 but is non-responsive to CIF2 (**Figure 2F**), other examples include *WRKY30*, *LAC1*, *PER52* (**Table S1, Movie S3).** Some reporters respond equally well to either flg22 or CIF2 in *2-in-Endo,* many showing non-cell autonomous responses in surrounding root cell-types (e.g., *WRKY41*, *PER71*, *ERF105*) (**Table S1)**.

Intriguingly, our dataset revealed that the top 10% most responsive genes (by fold change) in early timepoints are common to both pathways, whereas genes responding most weakly (but significantly) exhibited more pathway specificity (**Figure S2E**). We speculate that “fast and strong” responses are often regulated by the common signalling modules of the two pathways.

### The transcription factor MYB36 controls SGN3-specific transcriptional responses

Our transcriptional analyses suggested that the transcription factor MYB36, reported to be a key regulator of endodermis differentiation,^19, 31^ also acts as a hub to regulate SGN3-specific responses. We found that genes regulated by MYB36 (downregulated in *myb36* compared to wild-type^31^), tend to respond specifically to CIF2 (**Figure 2E**). While the scale of induction by CIF2 is relatively weak (log_2_FC<2), many of these genes are specifically expressed in the endodermis and are known to contribute to CS integrity. For example, expression of *CASP* genes (*CASP1/2/3/4*) is crucial for establishing functional CSD,^34, 36^ while peroxidases such as PER9 and PER64^34^ mediate CS lignification,^37^ and the dirigent protein ESB1^35^ is essential for both CSD formation and lignification.^38^ These genes might therefore carry out SGN3-specific functions and their non-responsiveness to flg22 can straightforwardly explain the inability of flg22 to promote CS domain fusion.^19,313134,363738^

CIF2-induced CASP domain fusion requires *de novo* protein synthesis.^20^ We hypothesize that MYB36-mediated transcriptional responses upon SGN3 stimulation are at the core of CASP domain fusion, and that the FLS2 pathway is unable to activate MYB36 and thus all the crucial genes required for CASP domain fusion. Consequently, although the SGN3 pathway would function independently of MYB36 for ectopic lignification, the CASP & CS domain fusion absolutely requires MYB36. Conversely, MYB36 is partially functional in *sgn3* and *cif1 cif2*, but only mediates formation of aligned, but discontinuous CASP domain islands. SGN3 would therefore drive domain fusion by boosting the activity of MYB36.

### Predicted MAP kinase phosphosites in MYB36 are important for CS domain fusion downstream of the SGN3 pathway

MYB36, predicted to be phosphorylated by MPKs based on an *in vitro* protein microarray,^39^ contains three putative MPK phosphorylation motifs (**Figure S3A**). We tested the necessity and sufficiency of MYB36 phosphorylation at these sites for CS domain fusion by generating a phospho-null variant MYB36^AAA^ by substituting the three serines with alanines, and a phosphomimic variant MYB36^DDD^ by mutating them to aspartic acids.

The untagged MYB36^AAA^, introduced in *myb36* mutant, can restore CASP1 expression and accumulation in the endodermis, but often failed to complement CASP1 domain fusion (**Figure 3A, Table S2**). This suggests that MYB36^AAA^, while active, is less functional than either the wild-type MYB36^WT^ or the phosphomimic MYB36^DDD^, both of which fully complement CASP1 domain fusion (**Figure 3A, Table S2**). The fusion defects exhibited by MYB36^AAA^ can be described as a band of “bubbles” roughly aligned with the endodermis central plane (**Figure 3B**). These CASP1 bubbles consist of many smaller disconnected patches, in comparison to the larger, more connected domains caused by the loss of SGN3 signalling in *sgn3* or *cif1cif2* (**Figure 3A, B, C, S3C**). Further phenotyping of the MYB36 variants showed that, as expected, the ability to fuse CASP1 domains correlates with the formation of a continuous CS lignin (**Figure 3B)** and an early formation of an extracellular diffusion barrier (**Figure 3D**).

Importantly, we found that the domain fusion defect in *myb36* MYB36^AAA^ is resistant to external CIF2 application (**Figure 3B, C, S3C, S3D**), suggesting that CIF2-induced domain fusion is impaired by the absence of functional phosphosites in MYB36 (**Figure 3F** middle). By contrast, CIF2-induced ectopic lignification is unaffected by MYB36^AAA^. The presence of ectopic lignin in *myb36* MYB36^AAA^, indicative of CS barrier defects that allows CIF peptide leakage, is further enhanced by external CIF2 application (**Figure 3B**). This phenotype is consistent with the delayed PI block in *myb36* MYB36^AAA^, similar to the untransformed *myb36* (**Figure 3D)**. Our data strongly suggest that the SGN3 pathway operates through multiple branches, one requiring MYB36 activation for CASP domain fusion, likely via phosphorylation, and another triggering the compensatory ectopic lignification independent of CASP accumulation and MYB36 involvement (**Figure 3F** left). Specificity could thus be attributed to the FLS2 pathway’s inability to activate MYB36.

**Figure 3.**
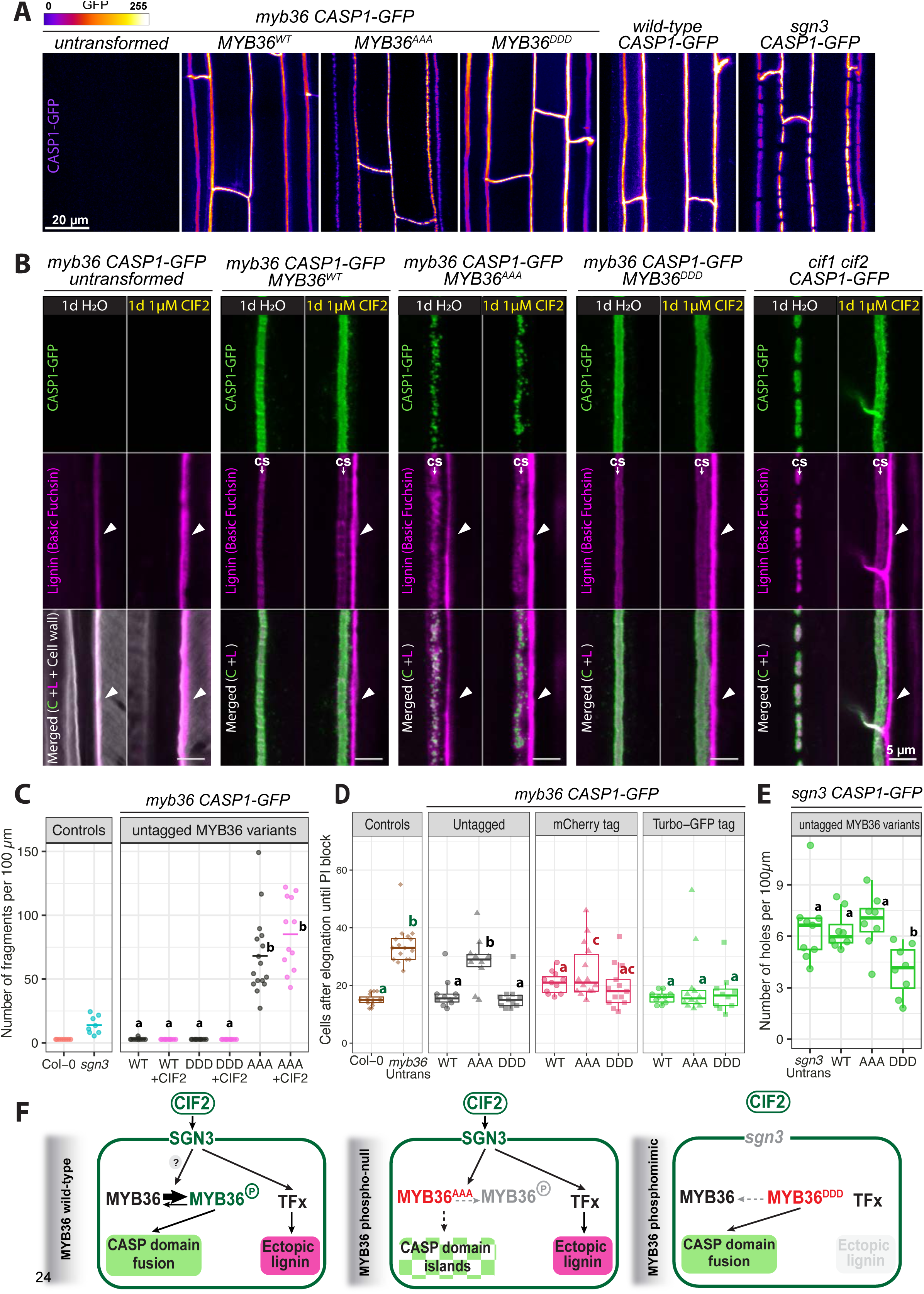
Predicted phosphosites of MYB36 are crucial for SGN-dependent CASP and CS domain fusion. (A) CASP1-GFP (gradient intensity) in *myb36* CASP1-GFP transformed with untagged MYB36 genomic DNA constructs: MYB36^WT^, MYB36^AAA^ (S18A, S146A, S169A), MYB36^DDD^ (S18D, S146D, S169D). T2 roots at mature CS stage (16-17^th^ cell) are shown. Untransformed *myb36*, *wild-type* and *sgn3* withCASP1-GFP are shown as controls. T1 phenotype distribution see Table S2. (B) CS & CASP1 domain fusion in *myb36* transformed with untagged MYB36 variants after 1d 1µM CIF2 or H_2_O treatment. Endodermal cell surface views at mature CS stage (16-17^th^ cell) show CASP1-GFP (green), Lignin (magenta) and overlaps of the two channels (Merged). Arrowheads: ectopic lignin on cortex-facing side. (C) Quantification of CASP1 domain properties: number of CASP1-GFP fragments per 100 µm in surface views. *myb36 CASP1-GFP* transformed with untagged MYB36^WT^, MYB36^AAA^, MYB36^DDD^ (WT, AAA, DDD) are treated with 1d H_2_O (grey) and 1d 1µM CIF2 (pink) (n≥10). Fragment size see Figure S3C. (D) PI penetration assay quantifies CS barrier function of *myb36* CASP1-GFP transformed with MYB36 constructs (WT, AAA, DDD), untagged or with C-terminal mCherry or Turbo-GFP tag (T1 roots, n≥10). Multiple comparisons are done separately for each tag group, always including 2 controls and 3 lines per tag group. (E) Quantification of CS discontinuity of *sgn3 CASP1-GFP* transformed with untagged MYB36^WT^, MYB36^AAA^ and MYB36^DDD^ (T1 and untransformed; n≥8). (F) Schematic models. SGN3 signalling acts through MYB36, possibly by boosting phosphorylation, to promote CASP domain fusion. Ectopic lignin is induced by SGN3 signalling independently of MYB36; MYB36^AAA^ is sufficiently active to induce CASP domain islands. MYB36^AAA^ cannot mediate CIF2-stimulated domain fusion, but can mediate CIF2-stimulated ectopic lignin acts via unknown transcription factor (TFx); MYB36^DDD^ is able to enhance CASP domain fusion without the SGN3 pathway. All data from roots at mature CS stage (16-17^th^ cell). For (C)(D)(E), groups with the same letter are not significantly different (P<0.05, one-way ANOVA and Tukey HSD test). See also Figure S3, Table S2.

We also found that different C-terminal tags affect the functionality of MYB36^AAA^, with large tags such as 6xHis3xFLAG-TurboID-GFP (Turbo-GFP, 847 aa) and mCherry (406 aa) apparently enhancing functionality of all MYB36 variants compared to their untagged version (**Figure 3D, Table S2)**. This might be due to enhanced stability of the protein fusions. Nevertheless, lines carrying MYB36^AAA^-mCherry, despite similar accumulation levels to MYB36^WT^-mCherry or MYB36^DDD^-mCherry, are still unable to fully complement CASP domain fusion in *myb36,* unlike the other variants (**Figure S3B**).

The phosphomimic MYB36^DDD^ could fully complement CASP domain fusion and CS barrier function in *myb36*, comparable to MYB36^WT^ (**Figure 3A-D**). Interestingly, the untagged MYB36^DDD^ shows gain-of-function activity, improving CASP domain fusion in *sgn3,* whereas MYB36^AAA^ could not (**Figure 3E, F** right). This further supports that the predicted MPK phosphosites in MYB36 are essential for CASP domain fusion induced by the SGN3 pathway. Furthermore, MYB36 protein accumulation and stability appear to be important determinants of its activity within the SGN3 pathway.

### Multiple MPKs contribute to CS integrity downstream of the SGN3 pathway

MPKs are key regulators of transcription factors, and our data indicate their potential in directly regulating MYB36 activity. Several MPKs are reported to be activated by both SGN3^20^ and FLS2 pathways.^26, 40^ In *2-in-Endo* seedling roots, we confirmed that both pathways can activate MPK3 and MPK6 in the endodermis upon flg22 or CIF2 treatments (**Figure 4A**). To obtain a more global picture of MPK activities in the endodermis, we first generated an expression atlas for all 20 MPKs in Arabidopsis (**Table S3**). We mapped the expression patterns of all MPKs by generating and analysing individual transcriptional fluorescent marker lines. Overall, we observed that MPKs are widely expressed across different tissues (**Table S3**), fitting with their roles in a wide range of cellular functions. We found that at least 12 MPKs are expressed in the endodermis and could thus be involved downstream of the SGN3 and FLS2 pathways and mediate signalling specificity (**Figure S4B, C**). Integrating our atlas with RNAseq^31^ and Translating Ribosomal Affinity Purification (TRAP) data enriched in the endodermal cell file (**Figure S4A**), we homed in on 8 candidates (MPK2/3/6/9/15/16/17/19). None of the tested *mpk* single mutants show strong defects in CS barrier function or CASP1 domain fusion (**Figure 4B, S4G**). We nevertheless observed a slight delay in forming a diffusion barrier in *mpk2* (**Figure 4B**), but no obvious discontinuities persisting at maturity were observed in *mpk2* CASP1-GFP (**Figure S4G**).

**Figure 4.**
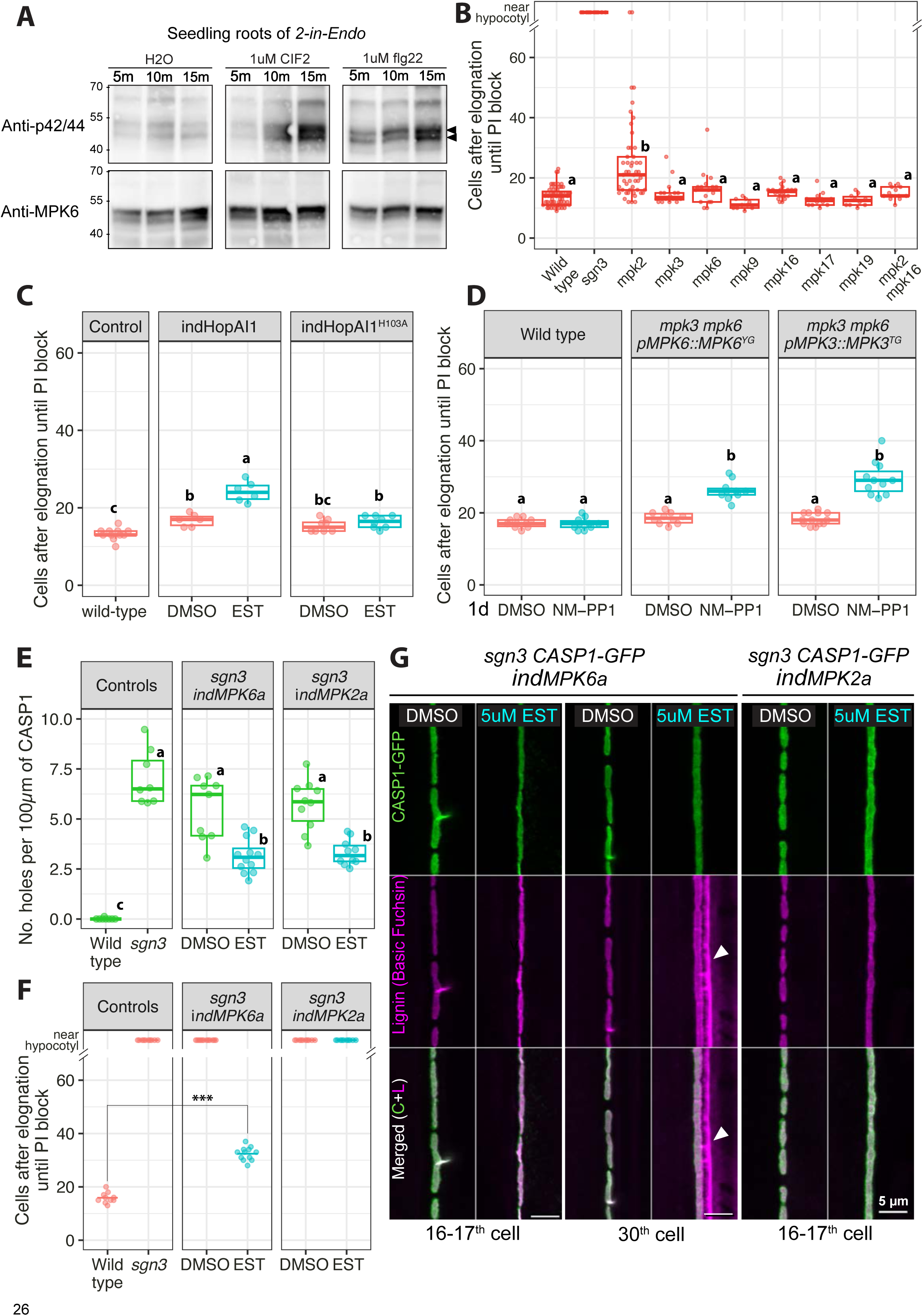
Multiple MPKs contribute to CS integrity downstream of the SGN3 pathway. (A) Immunoblots (IB) against p42/44 (phosphorylated MAPKs) indicate sizes corresponding to phosphorylated MPK6 (upper) and MPK3 (lower). *2-in-Endo* seedling roots treated with H_2_O, lµM CIF2 or flg22 for 5, 10, 15 mins. Loading controls: IBs with anti-MPK6 antibody. (B) PI penetration assay quantifies CS barrier function in various *mpk* mutants (10≤n≤49) compared with wild-type (n=63) and *sgn3* (n=10). Data from multiple independent experiments are combined. Unbalanced Tukey HSD test for unequal replication. (C) PI penetration assay with endodermal suppression of multiple MPK activities by inducing HopAI1 vs inactive HopAI^H103A^ in wild-type (n ≥ 6). Endodermis-specific estradiol (EST) inducible promoter: pLTPG15::XVE>>pLexA::HopAI1 (indHopAI1). (D) PI penetration assay upon conditional loss-of-function of both MPK3 and MPK6 (n≥10). pMPK6::MPK6^YG^ or pMPK3::MPK3^TG^ are sensitive to NM-PP1 in *mpk3 mpk6* background. 4d-old seedlings roots transferred to 5µM NM-PP1 or DMSO for 1d. (E) Quantification of CS discontinuity upon endodermal induction of constitutively active MPK6a or MPK2a in *sgn3 CASP1-GFP* (n≥8). (F) PI penetration assay upon endodermal induction of MPK6a or MPK2a in *sgn3 CASP1-GFP.* Student’s t-test compares indMPK6a to the wild-type (***P<0.001). (G) CS lignin & CASP1 phenotypes upon endodermal induction of MPK6a or MPK2a in *sgn3 CASP1-GFP* at 16-17^th^ and 30^th^ endodermal cells after onset of elongation. Surface views show CASP1-GFP (green) and Lignin (magenta). Arrowheads: ectopic lignin on cortex-facing side. For (B)(C)(D)(E), groups with the same letter are not significantly different (P<0.05, one-way ANOVA and Tukey HSD). *“*ind” designates “pLTPG15::XVE>>pLexA::”. For EST induction, all seedlings were grown for 5d on 5 µM EST or DMSO non-induced control before analysis. See also Figure S4, Table S3.

Because MPKs are involved in many processes, dissecting their function without pleiotropic effects in higher order mutants is challenging. To navigate this, we used HopAI1, a *Pseudomonas syringae* effector that specifically deactivates multiple MPKs, including MPK3/4/6, to suppress host immune signalling.^41–43^ To exert precise spatiotemporal control of MPK suppression via HopAI1, we used the estradiol (EST) inducible promoter active in endodermis: *pLTPG15::XVE>>pLexA::HopAI1* (hereafter indHopAI1). HopAI1 induction in wild-type plants leads to a delayed barrier formation and some ectopic lignin suggesting that MPK activities are required for CS functionality, but no phenotype is observed upon induction of an inactive HopAI1^H103A^ mutant (**Figure 4C, S4D**). However, HopAI1 induction in the endodermis does not seem to affect CASP domain fusion at our confocal resolution (**Figure S4D**). This suggests that either HopAI1 does not target MPKs required for CASP domain fusion, or that is does not sufficiently interfere with their activity.

As *mpk3 mpk6* double mutants are reported to be embryo-lethal,^44^ we used the conditional loss-of-function lines expressing *pMPK3::MPK3^TG^* or *pMPK6::MPK6^YG^* in an *mpk3 mpk6* background.^45, 46^ MPK3^TG^ and MPK6^YG^ are specifically sensitive to NM-PP1, an analogue of the general kinase inhibitor PP1. We found that one-day inhibitor treatment was sufficient to delay the PI block without causing other developmental defects (**Figure 4D**). In contrast, a one-day NM-PP1 treatment on wild-type causes neither developmental nor barrier defects (**Figure 4D**).

We also use a gain-of-function approach to investigate if constitutively active MPK6^D218G/E222A^ or MPK2^Q188G/E192A^, designated MPK6a and MPK2a, can alleviate CS defects in *sgn3*. We found that expressing MPK6a or MPK2a under an estradiol-inducible endodermis-specific promoter (ind) in *sgn3* can partially induce CASP1 domain fusion, leaving reduced discontinuities at maturity (**Figure 4E**). Induction of MPK6a, but not MPK2a partially restores the diffusion barrier in *sgn3* (**Figure 4F)**. This correlates with MPK6a activating ectopic ROS (**Figure S4F**) and ectopic lignin and suberin formation in *sgn3*, unlike MPK2a (**Figure 4G, S4E**). MPK6a appears to have a broader activity, affecting both the specific and common branches of the SGN3 pathway, while MPK2a contributes more specifically to CASP domain fusion.

Since MPK6a is sufficient to induce ectopic lignin in *sgn3*, we investigated whether MPK6 is necessary for SGN3-dependent ectopic lignification by testing if CIF2-induced ectopic lignin is affected in *mpk6*. We found that one-day CIF2 treatment could induce similarly strong ectopic lignin in *mpk6* and wild-type, as well as for *mpk3* (**Figure S4H**). This suggests that genes in addition to MPK6 must contribute to ectopic lignification downstream of the SGN3 pathway. Despite MPK2a and MPK6a contributing to domain growth and fusion, neither alone is sufficient to complete CS fusion. Therefore, a combination of different MPK activities appears to contribute to the three-branched outputs of the SGN3 pathway, some unique, some overlapping with the FLS2 pathway.

### Activation of individual MKKs in the endodermis produces distinct outputs, matching the common and specific outputs of SGN3 and FLS2 pathways

Contrasting the 80 MAPKKKs and 20 MAPKs, only 10 MKKs are found in Arabidopsis,^47, 48^ and it is unclear how much specific information can be transduced through this comparative bottleneck. MKKs tend to selectively activate certain MPK combinations,^39, 49^ suggesting that the activation of different MKKs may translate into distinct combinations of MPK activities, depending on the specific pathway triggered.

Given the relatively small number of MKKs, it is feasible to conduct a comprehensive examination of their activities in the endodermis. Since we did not find even weak phenotypes in any of the single and double mutants tested (**Figure S5K, M, N**), we decided to use our endodermal specific induction system and introduce all 10 constitutively active MKKs in the *sgn3* background (designated indMKK1a to indMKK10a). The 10 MKKs are classified into phylogenetic groups (A-D)^47^ ^43^(**Figure S5A**), and interestingly these groupings match strikingly well with the functional outcomes we observe in the endodermis.

For Group A/B members, namely MKK1/2/3/6, we found that induction of each individual MKKa in the *sgn3* endodermis can improve CASP1 and CS lignin fusion (**Figure 5A, B, S5J**), although none is sufficient to rescue the barrier defects of *sgn3* in the diffusion assay (**Figure 5C**). This can be attributed to the persistence of small gaps in the CS after induction (**Figure 5A**), coupled with the fact that these activated variants do not induce compensatory ectopic lignin (**Figure 5A, S5J**). Intriguingly, unlike MPK6a, which triggers early excess suberin, MKK1/2/3a induction suppresses suberisation in *sgn3* roots (**Figure S5H**). Our findings suggest that MKK1/2/3/6 are involved in transducing the SGN3-specific branch of CS domain fusion, which aligns nicely with the *in vitro* data of MKK1-MPK2 and MKK2-MPK2 phosphorylating MYB36.^39^ This fits our model, whereby the divergent outputs of SGN3 and FLS2 are defined by differential regulation of MYB36 via distinct MKK-MPK signalling modules (**Figure 7**).

**Figure 5.**
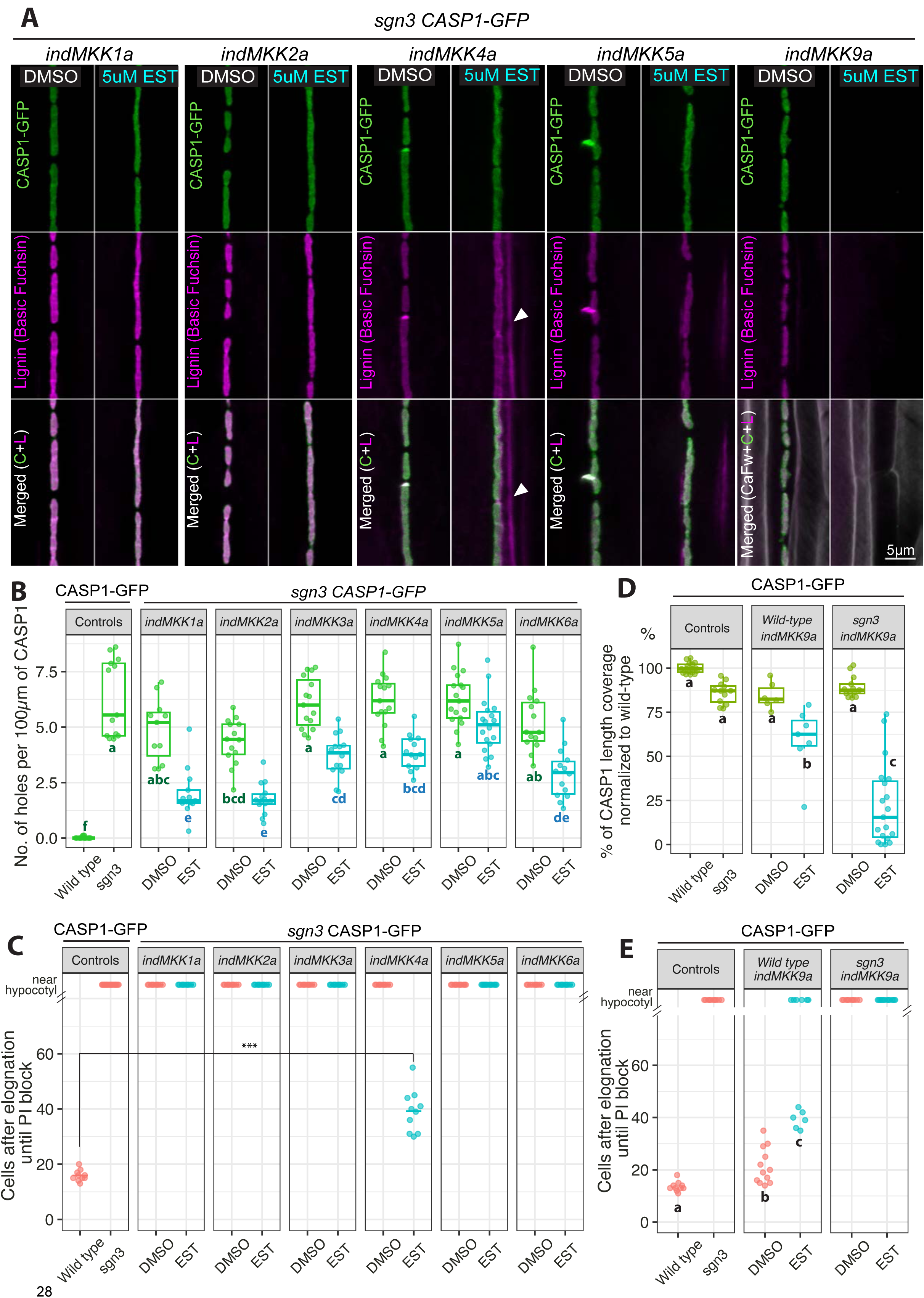
Activation of individual MKKs in the endodermis produces distinct outputs. (A) CASP1 & lignin phenotypes upon endodermal induction of constitutively active MKK1/2/4/5/9a in *sgn3 CASP1-GFP* at 16-17th cell. Endodermal cell surface views show CASP1-GFP (green), Lignin (magenta). Right-most panel: cellulosic cell walls stained with Calcofluor White (grey) to visualise endodermal cell edges. Arrowheads: ectopic lignin on cortex-facing side. (B) Quantification of CS continuity upon endodermal induction of MKK1/2/3/4/5/6a in *sgn3 CASP1-GFP* (n≥11). (C) PI penetration assay quantifies CS barrier function upon endodermal induction of MKK1/2/3/4/5/6a in *sgn3 CASP1-GFP* (n≥10). Student’s t-test compares indMKK4a and wild-type (*** P<0.001). (D) Quantification of CS coverage upon induction of MKK9a in *wild type* and *sgn3 CASP1-GFP* (n≥10). At mature CS stage, total CASP1 length of each overview image measured normalized to the average CASP1-GFP length in wild-type to calculate percentage of CS coverage for each data point. (E) PI penetration assay upon endodermal induction of MKK9a in *wild type CASP1-GFP*, and in *sgn3 CASP1-GFP* (n≥10). For (B)(D)(E), groups with the same letter are not significantly different (P < 0.05, one-way ANOVA and Tukey HSD). For EST induction, all seedlings were grown for 5d on 5µM EST or DMSO before analysis. For each indMKKa, two independent T2-line-data are combined in (B-E). See also Figure S5.

Group C members, MKK4 and MKK5 show broader activities. Induction of MKK4a can moderately improve domain fusion in *sgn3* (**Figure 5B**), but uniquely promotes ectopic lignin and ROS production, even in the absence of SGN3 signalling (**Figure 5A, S5L**). This ectopic lignin is sufficient to form a delayed diffusion barrier in contrast to no barrier with non-induced control in *indMKK4a sgn3* (**Figure 5C**). In addition, indMKK4a triggers early excess suberin accumulation (Fluorol yellow staining) and suberin synthesis (GPAT5 expression marker) in *sgn3* (**Figure S5H, I**). These data suggest that MKK4a mediates the compensatory branches of the SGN3 pathway (ectopic lignin & excess suberin), a function akin to MPK6a’s effects. Given that MKK4-MPK6 is activated by FLS2 for immune function,^26, 50^ this cascade could regulate the common downstream outputs of both SGN3 and FLS2 in the endodermis (**Figure 7**). MKK5, often considered functionally redundantly with MKK4 in immune signalling,^26^ has little effect on CASP1 domain fusion or ectopic lignification (**Figure 5A-C**), but does slightly enhance suberin synthesis upon induction in the *sgn3* endodermis (**Figure S5H, I**).

Among Group D members, MKK9 stands out as we found the others (MKK7/8/10) are either absent or only weakly expressed in the root endodermis (**Figure S5A**).^31^ Strikingly, EST induction of MKK9a in *sgn3* suppresses both CASP1 and CS lignin accumulation (**Figure 5A, D**). Electron microscopy confirms that MKK9a also inhibits ROS accumulation and membrane attachment, typical of a functional CS structure (**Figure S5L**). This suppression is specific to new growth and remains prominent in the presence of the SGN3 pathway in wild-type background (**Figure 5D, S5B**). In the wild-type, MKK9a suppresses CASP1 accumulation and domain fusion but does not suppress ectopic lignification (**Figure S5B**), aligning with the partial PI block phenotype (**Figure 5E**). MKK9a also has a potent inhibitory effect on suberin in *sgn3*, resulting in patchy or discontinuous suberization throughout the root (**Figure S5H, I**). Western blot analysis confirms the enhanced accumulations of MKK1/2/4/5/9a upon EST induction (**Figure S5F, G**). We observe MKK4a and MKK5a’s ability to enhance phosphorylation of MPKs, whereas induction of MKK1/2/9a do not trigger an increase in MPK phosphorylation (**Figure S5F, G**).

Despite being naturally low in the endodermis, inducing MKK8a and MKK10a in the *sgn3* endodermis improves CASP1 domain fusion (**Figure S5C**). However, neither MKK could alter the diffusion barrier defects (**Figure S5E**). Intriguingly, indMKK7a suppresses CASP1 accumulation in *sgn3*, mimicking indMKK9a (**Figure S5D**). This is consistent with previous reports that MKK7 shares functional similarity with MKK9 in ethylene signalling and stomatal development.^51, 52^

In summary, our systematic analysis of all 10 MKKs in a single cell-type demonstrates that activation of individual MKKs in the endodermis leads to clearly distinct output patterns. These patterns match both the common and specific outputs upon SGN3 and FLS2 stimulation in the endodermis. The Groups A/B (MKK1/2/3/6) are strong positive regulators contributing specifically to domain fusion; Group C, specifically MKK4, is the only one capable of promoting ectopic lignin and excess suberin, with MKK5 primarily influencing suberin. Finally, among Group D, MKK7 and MKK9 are strong negative regulators capable of suppressing CS development. These data strongly support our model that activation of distinct MKKs by SGN3 vs. FLS2 can explain the observed pathway specificities (**Figure 7**).

### Constitutively active MKK9 inhibits CS formation via suppression of MYB36

The specific impact of MKK9 activation on CASP1 accumulation and fusion suggests that it could be a powerful suppressor of MYB36 activity. We tested this by introducing MYB36^WT^, MYB36^AAA^ or MYB36^DDD^ tagged with TurboID-GFP (here labelled -GFP) into the *sgn3 indMKK9a* background, as this tag appeared to give the strongest activity (**Figure 3D, TableS2**). Regardless of the presence of MYB36^WT^-GFP or MYB36^AAA^-GFP, indMKK9a continues to suppress CASP1 accumulation in *sgn3* (**Figure 6A, B)**. In contrast, the phosphomimic MYB36^DDD^-GFP abolishes the inhibitory effect of indMKK9a on CASP1 accumulation (**Figure 6A, B**). This indicates that MKK9 suppression is conditioned by the phosphorylation status of MYB36.

**Figure 6.**
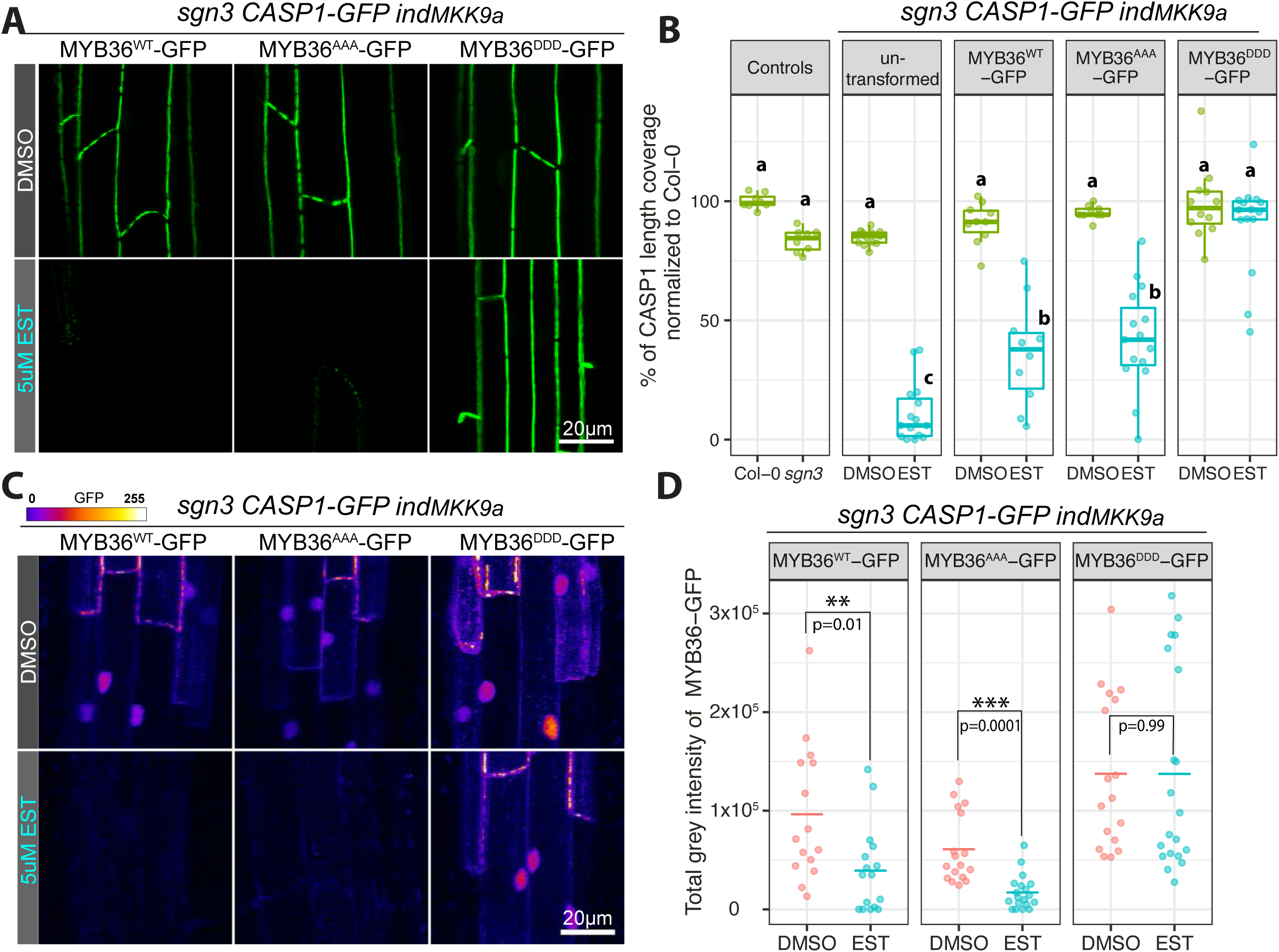
Constitutively active MKK9 inhibits CS formation via suppressing MYB36. (A) CASP1-GFP phenotypes of *sgn3 CASP1-GFP indMKK9a* (T3) transformed with Turbo-GFP tagged MYB36^WT^-GFP, MYB36^AAA^-GFP, MYB36^DDD^-GFP. T2 roots at mature CS stage (16-17^th^ cell) are shown. (B) Quantification of CS coverage upon induction of MKK9a *sgn3 CASP1-GFP* transformed with MYB36-GFP variants (T2, n≥8). Groups with the same letter are not significantly different (P<0.05, one-way ANOVA and Tukey HSD). (C) Stability of MYB36 variants in *sgn3 CASP1-GFP indMKK9a* background with or without EST induction. Nuclei signals of MYB36-GFP at 7-8^th^ cell (start of CASP1-GFP deposition) corresponding to (A) are shown. CASP1-GFP and MYB36-GFP signals can be distinguished based on different subcellular location. GFP is shown in gradient intensity. (D) Quantification of MYB36 variants stability as in (C), showing total grey intensity of nuclei signal (MYB36-GFP fluorescence) with or without EST induction at 7-8^th^ cell (T2, n≥15). Student’s t-tests compare EST and DMSO for each genotype. For EST induction, all seedlings were grown for 5d on 5µM EST or DMSO before analysis. See also Figure S6.

Furthermore, MKK9a induction reduces the accumulation of both MYB36^WT^-GFP and MYB36^AAA^-GFP in the endodermal nuclei (**Figure 6C, D**). The lower MYB36 levels correspond closely with the severity of CS suppression (**Figure 6A, C, S6**). Conversely, MYB36^DDD^-GFP levels were unaffected by MKK9a induction, with the strong MYB36^DDD^-GFP accumulation matching increased CASP1 coverage (**Figure 6**). Intriguingly, in *sgn3 indMKK9a* lines expressing high levels of MYB36^DDD^-GFP, we sometimes observed ectopic CASP1 patches that occur outside of the CSD (**Figure S6**), an outcome never observed for either MYB36^WT^-GFP or MYB36^AAA^-GFP lines. This further supports the idea that the phosphomimic variant of MYB36 possesses an enhanced ability to promote new CASP1 domain formation and growth, even in the context of *sgn3*. As this ectopic CASP1 phenotype of MYB36^DDD^-GFP mimics the hyperstimulation of the SGN3 pathway,^12^ we deduce that MYB36 is phosphorylated downstream of the SGN3 pathway.

Altogether, our results provide evidence that MKK9a’s suppression of CS directly correlates with phosphorylation status and stability of MYB36. MKK9 signalling appears to destabilize non-phosphorylated MYB36 to suppress its function. This supports our model that specific MYB36 phosphorylation and enhanced stability mediated by the SGN3 pathway is essential for its specific central output: precise domain fusion of the CASPs to form a continuous CS lignin barrier.

## Discussion

### Signalling specificity is maintained at single cell level

The complexity and intricacy of cellular signalling are increasingly recognized in plants.^53, 54^ Understanding the signalling dynamics at the single-cell level is fundamental for delineating how overall organismal responses are shaped. Despite this recognition, unravelling signalling specificity within a single cell remains challenging, owing to the difficulties in both stimulating and observing pathway read-outs at single cell resolution. Our study utilized the unique characteristics of the SGN3 pathway, enabling precise and quantifiable readouts in a non-dividing, easily observable cell.

Tissue specificity greatly influences signal interpretation and response by providing a contextual signalling landscape to a cell. Indeed, overexpression and ectopic expression of receptors can disrupt this signalling landscape and perturb output specificity in both plants and animal cells.^55^ Yet, if tissue compartmentalization were the primary insulating mechanism, any appropriately expressed foreign receptor/ligand could replace the native pathway.

By establishing a system that allows stimulation of both the FLS2 and SGN3 receptor pathways in the same cell type, we demonstrated that the FLS2 pathway in the endodermis could not substitute the SGN3 pathway in establishing a mature and continuous Casparian strip (**Figure 1**). Importantly, our *2-in-Endo* system reflects an endogenous signalling as the endodermis inherently has the capacity for FLS2 immune signalling, which is evident when endogenous FLS2 receptor level in the endodermis increases upon mechanical damage.^30^ As such, our study underlines the importance of examining signalling specificity beyond the constraints of tissue specificity and compartmentalization.

### Signalling pathways “copy and adapt” to achieve specialization

In the same cell, a simplistic approach to achieve specificity would be to have dedicated sets of receptors, signalling components and transcription factors that are insulated between one pathway and another in the same cell. However, evolution appears to have favoured a more elegant solution — reusing key signalling components from diverse processes, a common theme seen among multicellular organisms.^56, 57^

In plants, despite the expansion of the receptor repertoire for detecting a growing number of specific inputs, intracellular signalling pathways share striking similarities in their structure and components. It is thus reasonable to believe that outputs of many pathways could share a common core. This is supported by the extensive interconnections among plant hormonal pathways^58^ and the substantial overlap frequently observed in transcriptomic responses to biotic and abiotic stresses.^59–61^ Our data expand this idea to a developmental and an immunity receptor kinase pathway. We found that strong stimulation of both FLS2 and SGN3 pathways can induce common ROS, lignin and suberin production (**Figure 1**). This is corroborated by transcriptional profiles, showing matching induction patterns for genes involved in lignin and suberin biosynthesis following either CIF2 or flg22 treatments (**Figure 2**). These commonalities raise an intriguing question: Could both pathways have originated from a shared signalling pathway module, having diverged over time to perceive distinct inputs while maintaining partially overlapping outputs? This scenario fits nicely with the prevalent view of plant evolution being driven by cycles of gene duplication and neo-functionalization.^62^ For instance, once the plant developed a mechanism for ROS production during bacterial defence, it may have been evolutionarily expedient to adapt these responses for more precise developmental purposes, leading to the emergence of the primary function of the present-day SGN3 pathway.

Intriguingly, our comparative RNAseq analysis revealed a significant overlap among the genes most strongly responsive during early timepoints across the two pathways, implying a shared, “fast and strong” response. Yet, despite this remarkable overlap, our study clearly shows that an immune receptor kinase, even when strongly expressed in the right place and time, cannot replicate the central function of a developmental pathway.

### Combinatorial activation and cross-pathway inhibition of MAPK cascades encode specific signalling outcomes

Plant MPK cascades exhibit a complex, web-like structure, contrasting the simpler, more linear pathways found in yeast and mammals.^57, 63, 64^ The expansion of MPK cascade elements in plants has enhanced their capacity to form numerous combinations of MPKKK-MKK-MPK, allowing adaptability in signal transmission.^49, 57^ Our understanding of the plant MPK network interaction and activation has been advanced through global analyses, predominantly relying on *in vitro* methods.^39, 52, 65, 66^ However, the mechanisms that control specificity within these cascades remain largely unknown, in part due to a lack of extensive validation in plants, especially at the cellular level.

MPK3 and MPK6 are known to be activated by numerous signalling pathways associated with diverse cellular functions.^4, 67^ Similarly, MPK1/2/7/14 are activated by various stress pathways.^68, 69^ This wide-ranging activation make it challenging to understand how MPKs mediate specific cellular responses. It is often hypothesized that specificity arise from cell- or tissue-specific expression of MPKs or their target substrates. However, our reporter expression atlas (**Table S3**) and previous tissue-wide expression analyses indicate that MPKs are broadly expressed across tissues.^70, 71^ Our *2-in-Endo* line ensures that the downstream MPKs activated by both FLS2 and SGN3 pathways would encounter an identical set of target proteins. Based on our data, the specific activity of SGN3 can be attributed to the differential activation of MYB36, shown to be an MPK target *in vitro*.^39^ Our data strongly suggests this MPK phosphorylation is crucial for driving CASP and CS domain fusion downstream of the SGN3 pathway (**Figure 3**). Other endodermis expressed MYB transcription factors, such as MYB15 and MYB41 involved in lignin and suberin production respectively,^72, 73^ are also reported to be phosphorylated by MPKs^74, 75^ and might be mediators of the pathway responses common to FLS2 and SGN3, i.e., ectopic lignin and excess suberin production (**Figure 7**).

**Figure 7.**
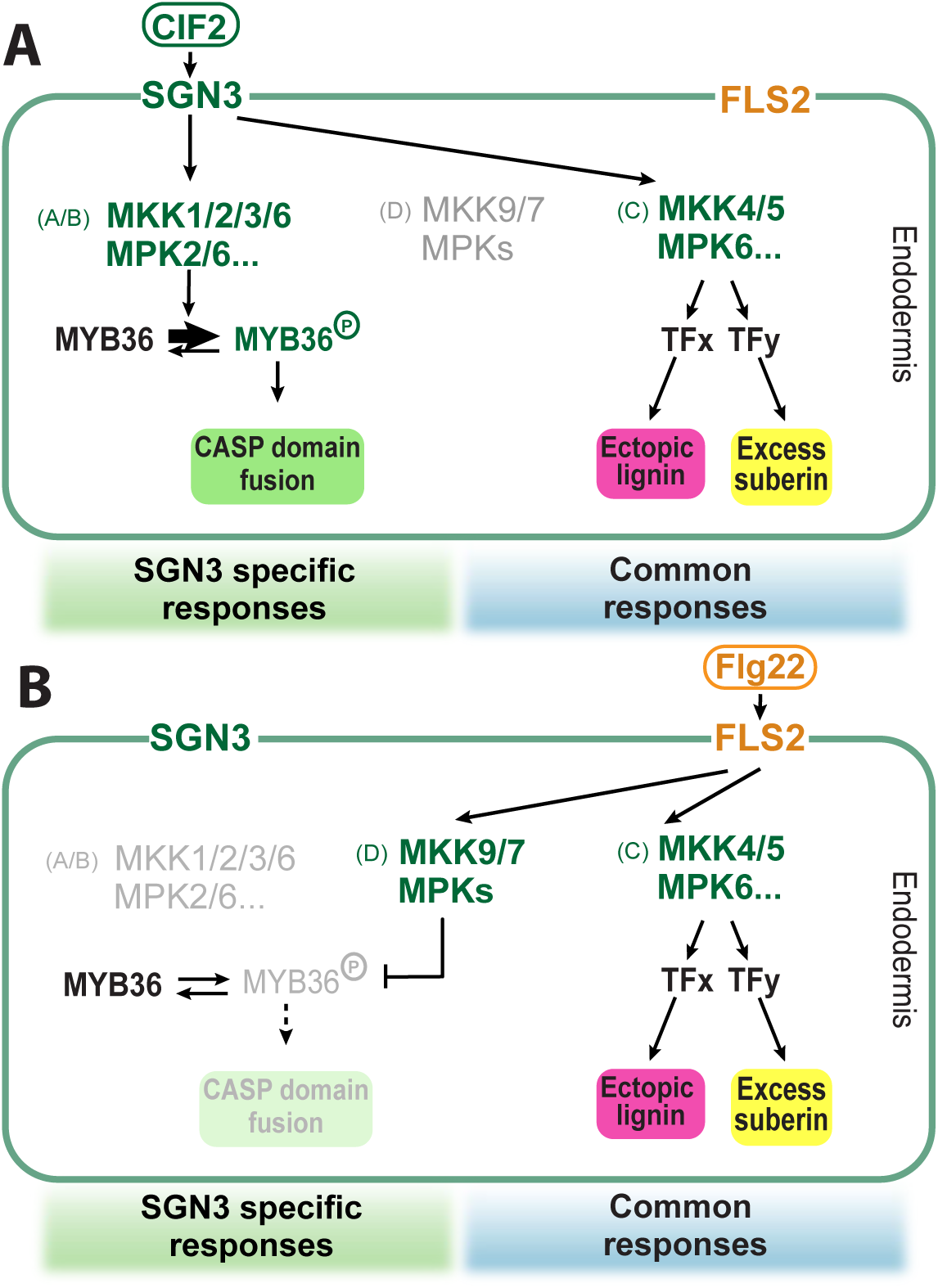
Current Model of two receptor pathways in a single cell type. Conceptual model of mechanisms underlying specificity of SGN3 signalling (A) and FLS2 signalling (B) in the endodermis.

In plants, MKK-MPK interactions are known to be promiscuous, with one MKK capable of activating multiple MPKs and many MKKs can activate the same MPK.^49, 52^ This promiscuity, while complicating analyses of single mutants, might be highly relevant for generating signalling specificity: different MKKs display preferential activity towards certain MPKs,^39, 49^ allowing each MKK to generate a potentially unique pattern of MPK activation. Each MPK could receive inputs from different MKKs depending on the pathway involved, and individual MPKs could activate both common and unique substrates,^76^ suggesting an extensive interaction network. Given that many independent MKKK-MKK routes can activate both the same and distinct MPKs, we propose that combinatorial MPK activation patterns generate distinct “flavours” that can drive specific outputs.

The elegant, comprehensive tissue-specific manipulations of MKKs in the stomatal lineage have demonstrated overlapping roles for MKKs 4/5 and 7/9 during early stages, and unique activities of MKK7/9 in late stages.^51, 77, 78^ In addition, MKK4/5, but not MKK1/2/3/8/9, contributes to root meristem development.^79^ These analyses revealed that MKKs have acquired divergent functions over time along a complex developmental trajectory. By contrast, our simpler endodermal system uses a differentiating, non-dividing cell-type that allows precise receptor stimulation and read-outs. Our comprehensive analysis of all ten MKKs in this system now provides compelling evidence that each MKK can activate distinct outputs within a single cell type (**Figure 5**). These functional groupings align well with their phylogeny and can account for both the common and unique outputs of the SGN3 and FLS2 pathways (**Figure 7**).

Given the matching readouts, our data suggest that SGN3 pathway’s central function primarily activates MKK1/2/3/6 (Groups A/B) to ensure domain fusion during CS development. However, upon strong or prolonged stimulation, the SGN3 pathway likely boosts MKK4/5 (Group C) activity to promote ectopic lignin and excess suberin. The SGN3 pathway may also avoid activation of negative regulators of CS development like MKK9 and MKK7 (in Group D), allowing effective MYB36 activation. Consistent with MKK9’s recognized role in ethylene signalling^80, 81^ - and the known inhibitory effect of ethylene activation on suberin in the endodermis,^82^-our data underscore that constitutively active MKK9 strongly suppresses suberin accumulation. Interestingly, the MKK groups contributing positively to SGN3 pathway outputs appear to have contrasting effects on suberin accumulation: Group C promotes suberin accumulation, Group A/B members suppress it. Among all MKKs, only MKK4 could enhance ROS production and lignification, consistent with a reported role of MKK4 in promoting ROS.^83^ Conversely, MKK5 and MKK3, but not MKK4, have been reported to suppress H_2_O_2_ (ROS) during various stresses.^84–86^ This suggests that ROS homeostasis could be maintained by balancing the activities of different MKKs downstream of SGN3. Altogether, these findings suggest an MKK-mediated tuning mechanism that balances SGN3 pathway responses, favouring CS domain closure upon low and transient stimulation, and preventing pleiotropic ectopic lignin & suberin accumulation until stronger, persistent stimulations are reached.

Based on the aligning readouts, we hypothesize that the FLS2 pathway in the endodermis has a weaker ability to activate MKK1/2/3/6 (Groups A/B), but more potently activates MKK4/5 (Group C), leading to enhanced ectopic lignin and suberin. Importantly, the inferred inability of FLS2 to activate MYB36 may be reinforced by a potentially stronger activation of MKK7/9, which would actively suppress MYB36 activity. Previous analyses of the FLS2 pathway paint a more complex picture with at least two independent MPK cascades, MKKK3/5-MKK4/5-MPK3/6 and MEKK1-MKK1/2-MPK4.^26, 28, 87, 88^ Initially thought to negatively regulate immunity, the MEKK1-MKK1/MKK2-MPK4 cascade was later revealed to activate guarding resistance proteins when perturbed.^43, 89–91^ Moreover, MKK6 suppresses autoactivation of the same guarding mechanism,^92^ while MKK7 contributes to flg22-induced ROS burst.^93^ Finally, the FLS2 pathway intertwines closely with ethylene signalling,^94–96^ known to activate MKK1/3/9.^80, 81, 97^

We contend that single cell-type analyses performed in a whole-tissue context offer clearer insights into signalling pathways and that the complexities observed in previous studies could be due to read-outs relying on protoplast experiments and/or immunoprecipitation of total plant tissues. It is intriguing to speculate that single cell signalling may involve frequent switching between signalling states. For instance, a strong stimulation of one pathway could actively suppress another, allowing a cell to respond to diverse stimuli, but not necessarily simultaneously.

### Limitations of the study

This study provides conclusive evidence of divergent functional outputs of ten individual MKKs and two MPKs in a single cell-type, using a gain-of-function approach. However, to genetically elucidate the MKK-MPK routes that lead to these distinct outputs requires complementary loss-of-function analyses. The major barrier is the redundancies and pleiotropic effects that arise from broad perturbation of multiple MPKs and/or MKKs. Potential work-arounds, such as cell-type specific MPK inhibition by HopAI1 phosphothreonine lyase activity appears to be restricted to a subset of MPKs.^41–43^ The specific MPKs affected and their activity after HopAI1 induction in the endodermis remain to be determined. Future studies would benefit from testing additional approaches for precise tissue specific MPK knock-outs.

Our data supports a compelling model of differential phosphorylation of MYB36 downstream of FLS2 and SGN3 pathways, driving output specificity. However, we have been unable to obtain sufficient amounts of MYB36 from *2-in-Endo* to determine their phosphorylation states after peptide treatment, due to the small number of responsive endodermal cells in total root cells. *In planta* biochemical validation of MYB36 phosphorylation would require a cell-type specific enrichment & phosphoproteomics approach. This approach could also help confirm and further delineate the MKK-MPK network coding output specificity in the endodermis.

## Acknowledgements

We thank the Cellular Imaging Facility (CIF), Genomic Technologies Facility (GTF) and Electron Microscopy Facility (EMF) of the University of Lausanne and Nick Pullen for technical support. We thank Gwyneth Ingram for critically reading the paper. We thank Zhang Shuqun and Jean Colcombet for kindly sharing seed materials. We thank all Geldner lab members for sharing materials. Y.M. was supported by EMBO non-stipendiary long-term fellowship (ALTF 711-2018), FEBS Long-term Fellowship and by H2020-MSCA-IF-2018 (Grant number 846050).

## Author Contributions

Y.M. and N.G. conceived, designed, and coordinated the project. Y.M., I.F., J.N., J.P., J.D., D.D.B., A.E. performed all experimental work and data analysis. S.F., A.E. provided materials and were involved in the discussion of the work. Y.M. and N.G. wrote the manuscript with feedback from all co-authors.

---

**Figure S1.**
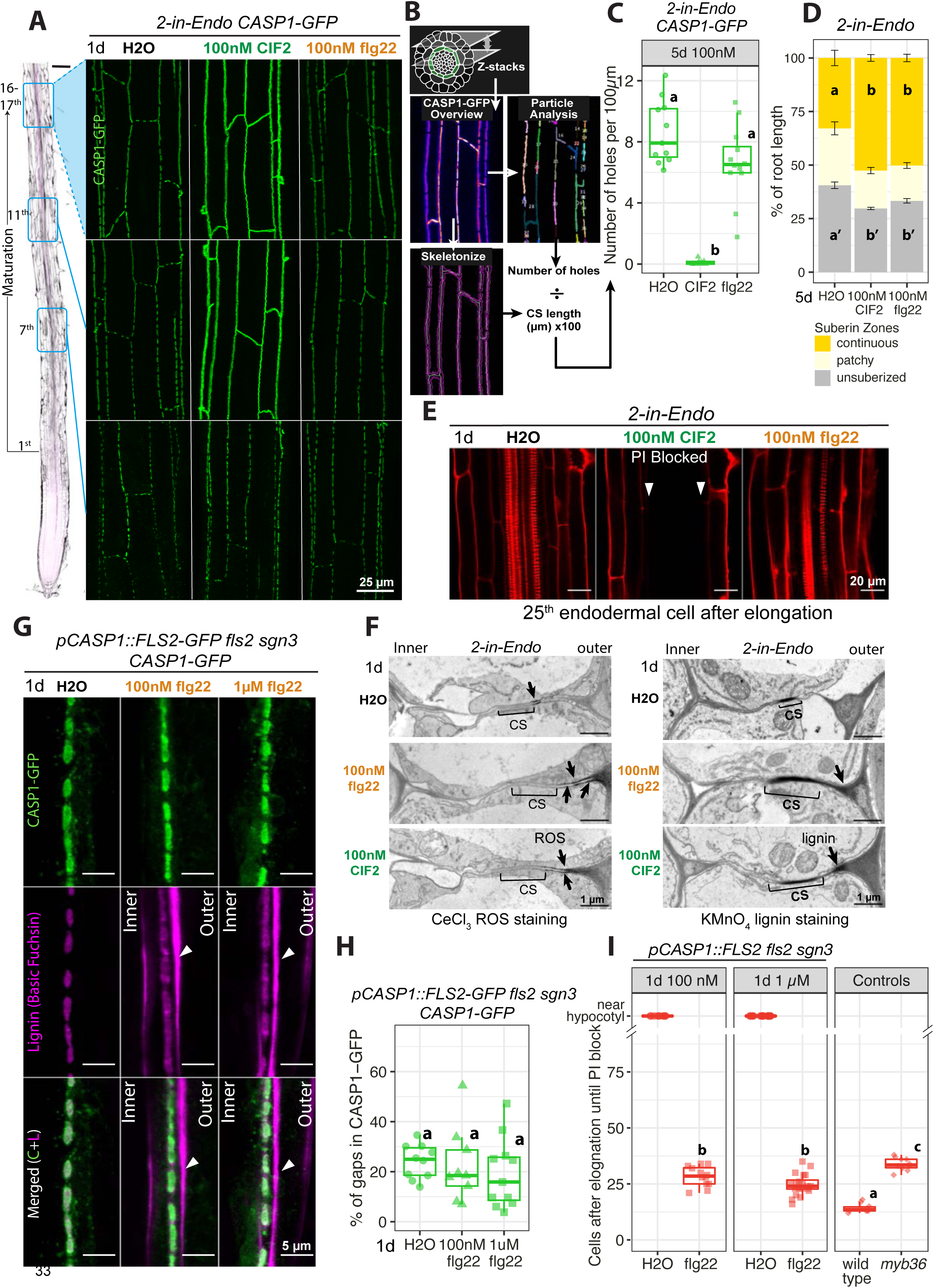
Endodermis-expressed FLS2 cannot functionally replace SGN3 to fuse Casparian Strip (CS) Domains. Related to Figure 1. (A) CASP1-GFP overviews of *2-in-Endo* after 1d treatments of H_2_O, 100 nM CIF2 or flg22 from initiation to maturation (7^th^, 11^th^, 16-17^th^ endodermal cells after onset of elongation). (B) Schematics showing steps to quantify CS discontinuity: 1. CASP1-GFP overviews: maximum projection of z-stacks capturing CS network in endodermal cell layer; 2. Particle Analysis (Fiji): count fragments and calculate corresponding number of holes; 3. Skeletonize (Fiji): convert area to lines and measure CS length in *µ*m; 4. Calculate “Number of holes per 100 µm” as in (C). (C) Quantification of CS discontinuity in *2-in-Endo* CASP1-GFP overviews at mature CS stage after constitutive (5d) treatments with H_2_O, 100 nM CIF2 or 100 nM flg22 (n≥8). (D) Suberization quantification of *2-in-Endo* after constitutive (5d) treatments with H_2_O, 100nM CIF2 or flg22. Suberin was stained with Fluorol Yellow and quantified along the root axis divided by three zones: unsuberized (near the root-tip); patchy; and continuous (near hypocotyl). Data presented as percentage of total root length (Mean±SE; n =10). Zones with the same letter are not significantly different among treatments (p<0.05, one-way ANOVA and Tukey test). (E) Diffusion barrier of *2-in-Endo* visualised by Propidium Iodide (PI, red) penetration at 25^th^ endodermal cell after onset of elongation. Arrowheads mark PI signal exclusion from the inner side of an endodermal cell (PI block). (F) Transmission Electron Microscopy (TEM) showing *in situ* H_2_O_2_ (ROS) detection with CeCl_3_ (left) and lignin detection with KMnO_4_ (right) at the CS of *2-in-Endo* after 1d H_2_O, 100 nM CIF2 or flg22 treatments. Micrographs obtained from sections ∼1. 5 mm away from the root tip. Brackets indicate CS: uniformly light grey cell wall areas with membrane attachment (left) and dense black staining (right). Arrowheads indicate ectopic accumulations of ROS (left) and lignin (right). (G) CS & CASP1 domain fusion phenotypes of *pCASP1::FLS2-GFP fls2 sgn3 CASP1-GFP* after 1d treatment with H_2_O, 100nM flg22 or 1µM flg22. Endodermal cell surface views at mature CS stage (16-17^th^ cell) show CASP1-GFP (green), lignin stained with Basic Fuchsin (magenta) and overlaps of the two channels (Merged). Arrowheads highlight ectopic lignin. “outer”, cortex-facing, “inner”, stele-facing endodermal surface. (H) Quantifications for (G) showing the percentage of gaps in CASP1-GFP *pCASP1::FLS2-GFP fls2 sgn3* after 1d treatments (n≥9). (I) PI penetration assay quantifies the CS barrier function of *pCASP1::FLS2-GFP fls2 sgn3* with 1d treatments of H_2_O and flg22 at 100nM and 1µM concentrations compared to controls. Roots with no barrier show PI penetration near root-hypocotyl junction, shown in the “near hypocotyl” category and are excluded from numerical statistical test (n≥10). For (C)(H)(I), groups with the same letter are not significantly different (P<0.05, one-way ANOVA and Tukey test). Unbalanced Tukey test used for unequal replication.

**Figure S2.**
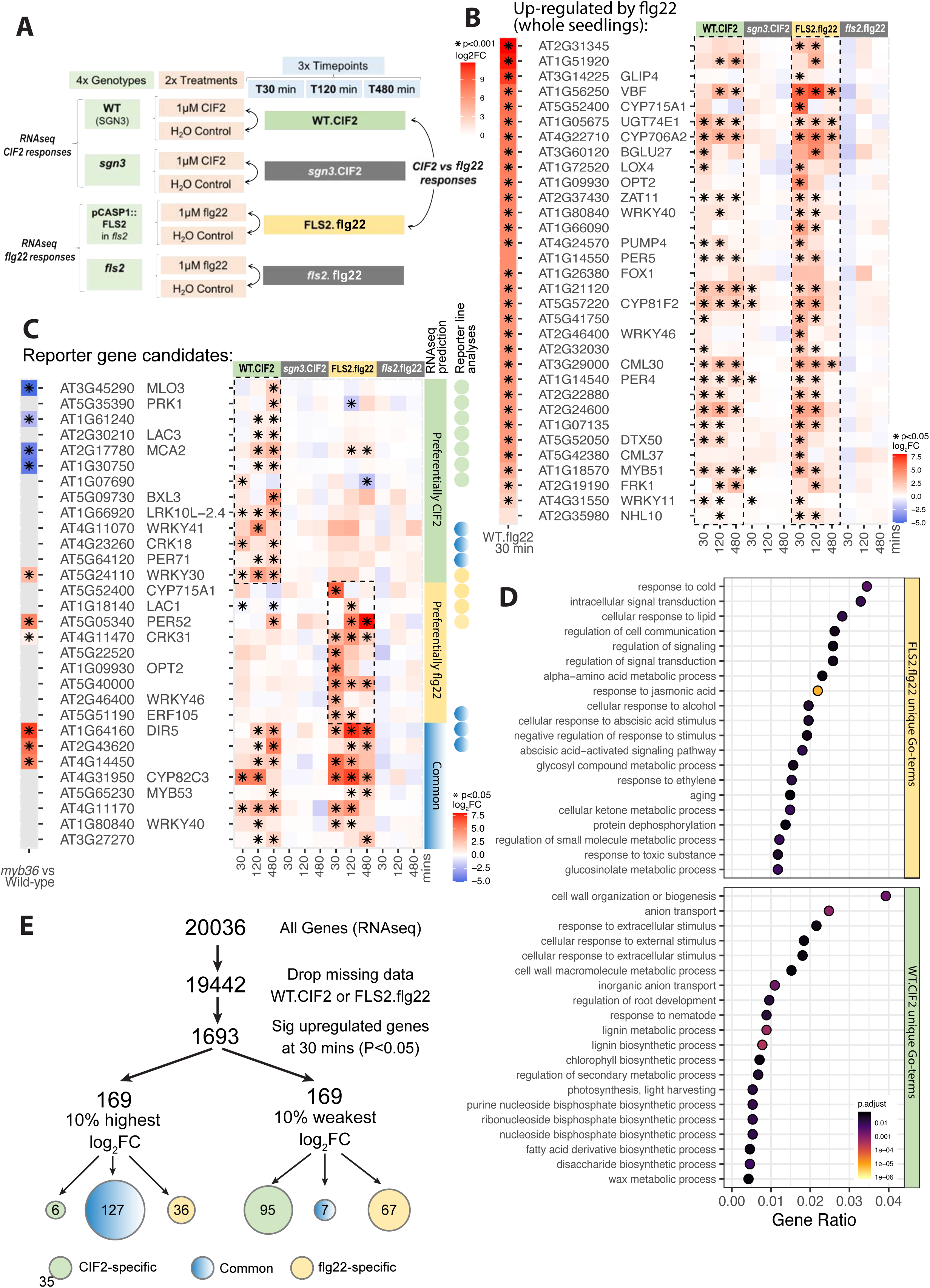
Endodermis-expressed FLS2 and SGN3 activate both common and pathway-specific transcriptional responses. Related to Figure 2. (A) Experimental setups for RNAseq analysis comparing CIF2 and flg22 responses in root endodermis. Transcriptional data for CIF2 responses are from Fujita et al.^1^ WT.CIF2, *sgn3*.CIF2: wild-type with functional SGN3 and *sgn3* treated with 1µM CIF2 vs H_2_O control. FLS2.flg22, *fls2*.flg22: pCASP1::FLS2-GFP *fls2* and *fls2* treated with 1µM flg22 vs H_2_O control. Fold change (log_2_FC) were calculated by comparing peptide (1µM flg22 or 1µM CIF2) to H_2_O treatment at each time point (30, 120, 480 mins). (B) Heatmaps of genes regulated by FLS2 in whole seedlings – significantly upregulated genes in flg22-treated wild-type (left) (*, P<0.001)^2^ – are also strongly induced by CIF2 (WT.CIF2) and flg22 (FLS2.flg22) in the endodermis (right) (*P<0.05). Colour code: fold change degree (log_2_FC). (C) Heatmaps of fold change for reporter genes. Candidates are predicted to be preferentially CIF2 (green) or flg22 (yellow) responsive, or equally responsive (blue) in the endodermis (*P<0.05). Reporter line analyses for selected genes are shown on the right (circles), with colour indicating observed pathway specificity (details see Table S1). Corresponding heatmap of fold-change in *myb36* compared to Wild-type (*P<0.001, NA in grey)^3^ shown on the left. Colour code: fold change degree (log_2_FC). (D) Gene ontology (GO) enrichment analysis showing enriched GO-terms unique to FLS2 and SGN3 pathways upon stimulation in the endodermis. Colour: GO-enrichment significance. Dot size: gene ratio (the gene counts in a GO category differentially regulated by a pathway divided by the total number of genes in that GO category). (E) Workflow to determine the pathway specificity of top 10% most responsive genes and least responsive (P<0.05, ranked by Log_2_FC) at 30 min. Genes that are significantly upregulated by one pathway but not the other are considered specific, genes that are significantly upregulated by both are considered common (Cut-off: Log_2_FC>1 and P<0.05).

**Figure S3.**
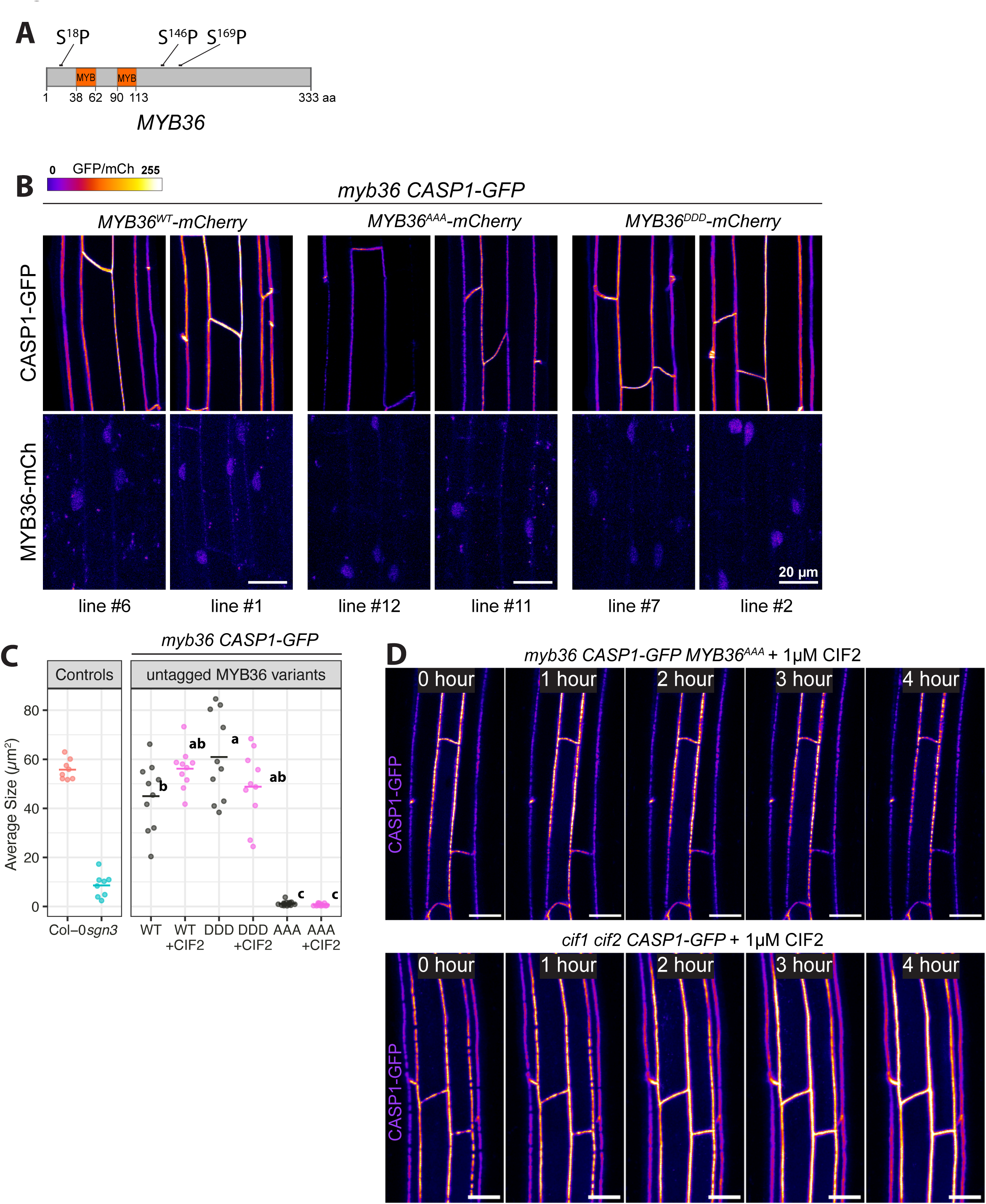
Predicted phosphosites of MYB36 are crucial for SGN-dependent CASP and CS domain fusion. Related to Figure 3. (A) *MYB36* protein diagram, with locations of three putative MPK phosphorylation sites (Serine Proline - SP) and two MYB-type HTH DNA binding domains (MYB). (B) CASP1-GFP phenotypes and MYB36 accumulations in *myb36* CASP1-GFP transformed with MYB36^WT^-mCherry, MYB36^AAA^-mCherry, MYB36^DDD^-mCherry. CASP1-GFP overviews (upper-panels) at 16-17^th^ cell and MYB36-mCherry nuclei signals (lower-panels) from the same lines at 7-8^th^ cells are shown for each genotype. GFP and mCherry in gradient intensity. (C) Quantification of CASP1 domain property: Average size of CASP1-GFP fragments in surface views, same data as Figure 3C. *myb36 CASP1-GFP* transformed with untagged MYB36^WT^, MYB36^AAA^, MYB36^DDD^ (WT, AAA, DDD) are treated with 1d H2O (grey) and 1µM CIF2 (pink) (n≥10). (D) CASP1-GFP time lapse images of *myb36 CASP1-GFP untagged MYB36^AAA^* (top) and *cif1 cfi2 CASP1-GFP ^A^* (bottom) treated with 1µM CIF2 from 0-4 hours. Maximum projections of roots at mature CS (16-17^th^ cell) are shown. Scale bars = 20µm

**Figure S4.**
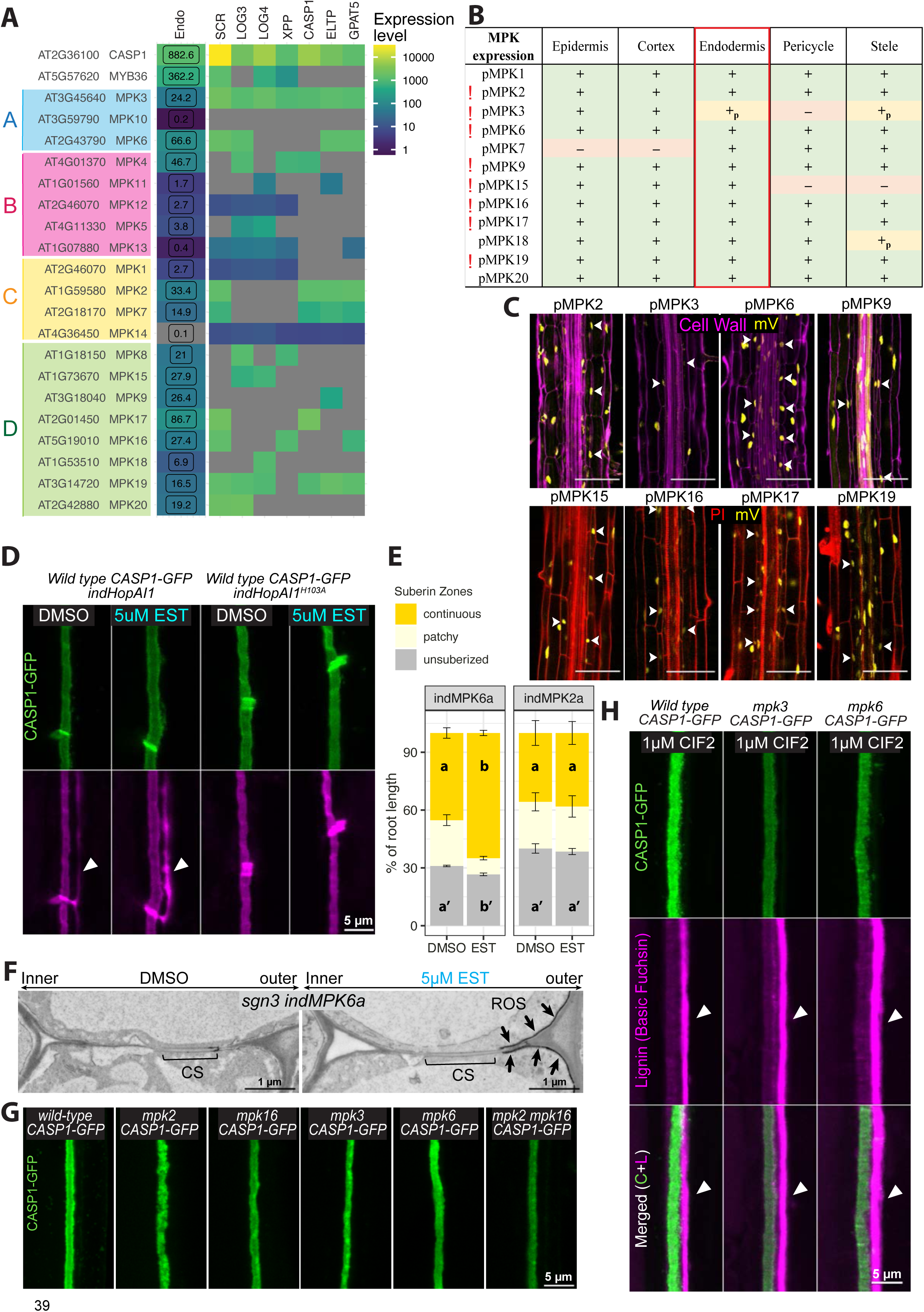
Multiple MPKs contribute to CS integrity downstream of the SGN3 pathway. Related to Figure 4. (A) MPK expression analysis in the endodermis. 1-20 MPKs from *Arabidopsis thaliana* are classified into 4 phylogenetic groups using full length sequences.^4^ Endodermis-enriched RNA-seq data^3^ (Endo) and Translating Ribosomal Affinity Purification (TRAP) analysis (Andersen & Vermeer personal communication) that detect mRNA transcripts in distinct compartments of endodermis and xylem poles using different promoters: SCR (Meristematic endodermis); LOG3 (Xylem pole protoxylem); LOG4 (Xylem pole endodermis); XPP (Xylem pole); CASP1 (Early unsuberized endodermis); ELTP or LTPG15 (The entire differentiated endodermis); GPAT5 (Suberized endodermis). CASP1 and MYB36 are used as references. (B) MPKs with endodermal expression around CS initiation (7-8^th^ cell). Analyses based on individual pMPK::NLS-3xmVenus lines (1 to 20) in wild-type. See Table S3 for full atlas analysis. Expression in all cells (green +), expression in some but not all cells (yellow +_p_), no expression in all cells (red –). “!” marks the selected MPKs for further analyses. (C) Examples of pMPK::NLS-3xmVenus reporter with endodermal signals during CS development. White arrows indicate endodermal nuclei. Cell walls are stained with Calcofluor white (magenta – upper) or with PI (red – lower). Scale bars = 100 µm. (D) CASP1-GFP phenotype with endodermal suppression of multiple MPK activities by inducing HopAI1 vs inactive HopAI^H103A^ in wild-type at 16-17^th^ cell. (E) Suberization quantification with or without MPK6a or MPK2a induction in *sgn3 CASP1-GFP*. Suberin stained with Fluorol Yellow and quantified along the root axis divided by three zones: unsuberized; patchy; and continuous. Data presented as percentage of total root length (Mean±SE; n≥10). Zones with the same letter are not significantly different (p<0.001, Student’s t-test). (F) TEM showing *in situ* H_2_O_2_ (ROS) detection with CeCl_3_ at the CS of *2-in-Endo* after 5d DMSO control or EST induction of MPK6a in *sgn3 CASP1-GFP*. Micrographs obtained from sections ∼3 mm from the root tip. Brackets indicate CS: uniformly light grey cell wall areas with membrane attachment. Arrowheads indicate accumulations of ROS. (G) CASP1 phenotypes of *mpk2, mpk16, mpk3, mpk6, mpk2 mpk16.* Surface views at 16-17^th^ cell are shown. (H) CS lignin & CASP1 phenotypes of *mpk3* and *mpk6* compared to wild-type, after 1d treatment of 1µM CIF2. Surface views at 16-17^th^ cell show CASP1-GFP (green), Lignin (magenta) and channel overlaps (Merged). Arrowheads highlight ectopic lignin on the cortex-facing side.

**Figure S5.**
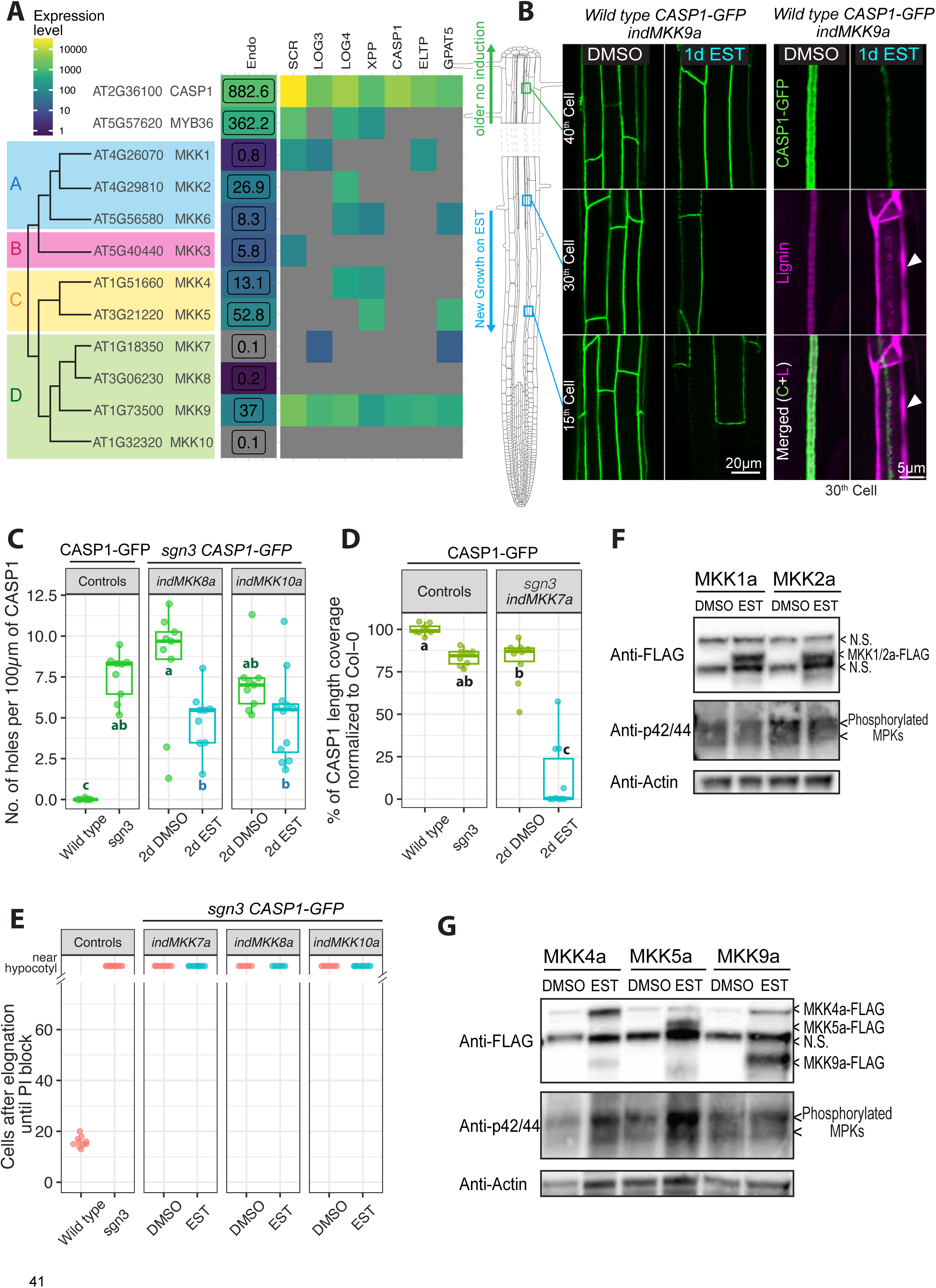
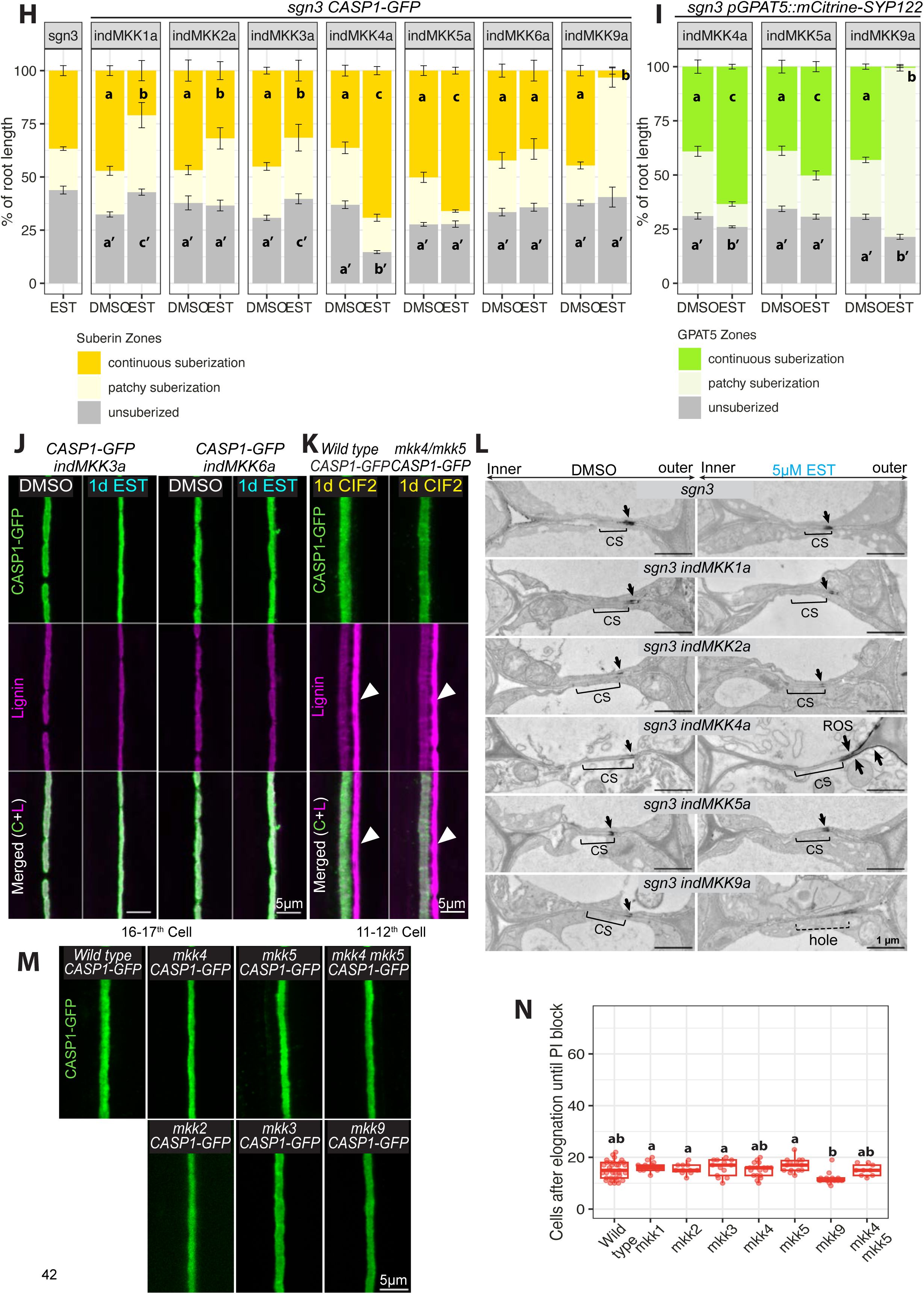
Activation of individual MKKs in the endodermis produces distinct outputs. Related to Figure 5. (A) MKK expression analysis in the endodermis. 1-10 MKKs from *Arabidopsis thaliana* are classified into 4 phylogenetic groups using full length sequence.^4^ Endodermis-enriched RNA-seq data ^3^ (Endo) and TRAP analysis (Andersen & Vermeer personal communication) that detect mRNA transcripts in distinct compartments of endodermis and xylem poles using different promoters: SCR (Meristematic endodermis); LOG3 (Xylem pole protoxylem); LOG4 (Xylem pole endodermis); XPP (Xylem pole); CASP1 (Early unsuberized endodermis); ELTP or LTPG15 (The entire differentiated endodermis); GPAT5 (Suberized endodermis). (B) CASP1 phenotypes after one-day EST induction of MKK9a in *wild-type CASP1-GFP* at new growth (15^th^ and 30^th^ cell -EST induction), and older growth (40^th^ cell - no induction). Right panels show zoomed surface views at 30^th^ cell for CASP1-GFP (green), Lignin (magenta) and channel overlaps (Merged). Arrowheads highlight ectopic lignin on the cortex-facing side. (C) Quantification of CS continuity: number of holes in CASP1-GFP per 100 µm upon endodermal induction of MKK8a and MKK10a in *sgn3 CASP1-GFP* (T1 roots, n ≥9). (D) Quantification of CS coverage upon induction of MKK7a in *sgn3 CASP1-GFP* (T1 roots, n =10). At mature CS stage, total CASP1 length of each overview image measured normalized to the average CASP1-GFP length in wild-type to calculate percentage of CS coverage for each data point. (E) PI penetration assay quantifies the CS barrier function upon endodermal induction of MKK7/8/10a in *sgn3 CASP1-GFP* (T1 roots, n≥10). (**F-G**) Immunoblots (IB) against anti-FLAG show accumulations of the C-terminally 3xFLAG tagged indMKK1a, indMKK2a (**F**) indMKK4a, indMKK5a, indMKK9a (**G**) in *sgn3 CASP1-GFP* background. T3 seedlings roots grown on DMSO or 5 µM EST were harvested. IBs against p42/44 (phosphorylated MPKs) indicate sizes corresponding to phosphorylated MPK6 (upper) and MPK3 (lower). N.S., non-specific bands. Loading controls: IBs with anti-Actin antibody. (H) Suberization quantification upon induction of MKK1/2/3/4/5/6/9a in *sgn3 CASP1-GFP*. Suberin stained with Fluorol Yellow and quantified along the root axis divided by three zones: unsuberized; patchy; and continuous. Data presented as percentage of total root length (Mean±SE; n≥10). (I) Suberin synthesis quantification upon induction of MKK4a, MKK5a and MKK9a in *sgn3* pGPAT5::mCitrine-SYP122 (SYP122 is a membrane marker). Suberin synthesis is marked by GPAT5 transcription. Data presented as percentage of total root length (Mean±SE; n≥10). For (H, I) Zones in each genotype with the same letter are not significantly different upon treatment (p<0.05, Student’s t-test for equal replicates, Welch t-test for unequal replicates). Zones with c/c’ are increased compared to a/a’, b/b’ decreased compared to a/a’. (J) CASP1 & lignin phenotypes upon endodermal induction of MKK3a and MKK6a in *sgn3 CASP1-GFP* at 16-17^th^ cell show CASP1-GFP (green), Lignin (magenta) and channel overlaps (Merged). Arrowheads highlight ectopic lignin on cortex-facing side. (K) CASP1 & lignin phenotypes of one-day 1µM CIF2 treated wild-type and *mkk4 mkk5 CASP1-GFP* at 11-12th Cell. Arrowheads highlight ectopic lignin on cortex-facing side. (L) TEM showing *in situ* H_2_O_2_ (ROS) detection with CeCl_3_ at the CS of *2-in-Endo* after 5d DMSO or EST induction of MKK1/2/3/4/5/9a in *sgn3 CASP1-GFP*. Micrographs obtained from sections ∼1.5-2 mm from the root tip. Brackets indicate CS: uniformly light grey cell wall areas with membrane attachment. Dashed bracket indicates the absence of an CS. Arrowheads indicate accumulations of ROS. (M) CASP1 phenotypes of *mkk2, mkk3, mkk4, mkk5, mkk9, mkk4 mkk5* at 16-17^th^ cell. (N) PI penetration assay quantifies the CS barrier function of *mkk1*, *mkk2, mkk3, mkk4, mkk5, mkk9, mkk4 mkk5* (n≥10). For (C)(D)(N), groups with the same letter are not significantly different (P<0.05, one-way ANOVA and Tukey HSD). For EST induction, all seedlings were grown for 5d on 5 µM EST or DMSO control before analysis unless indicated otherwise.

**Figure S6.**
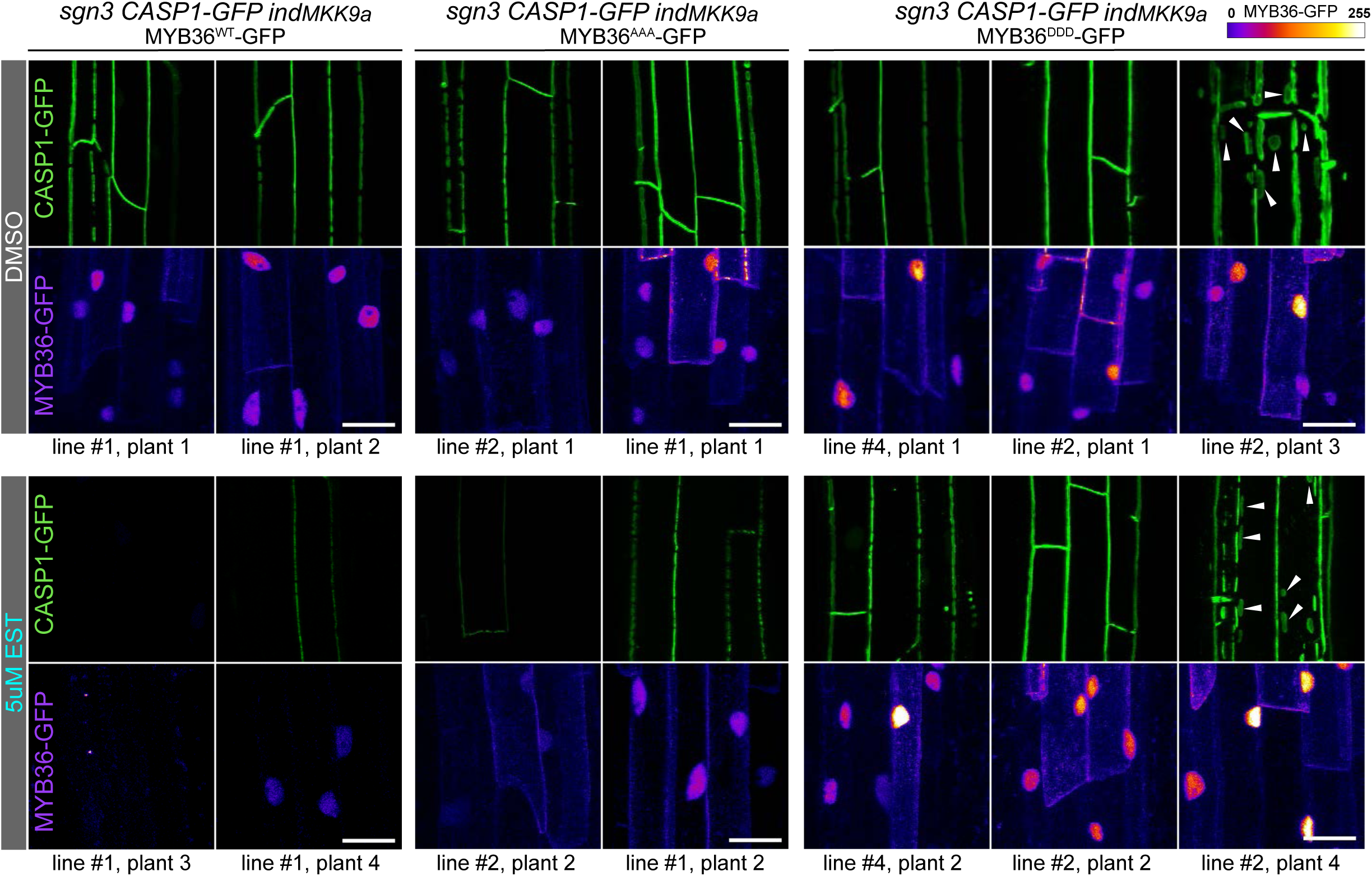
Constitutively active MKK9 inhibits CS formation via suppressing MYB36. Related to Figure 6. CASP1-GFP and corresponding MYB36 accumulations in *sgn3 CASP1-GFP indMKK9a* transformed with MYB36^WT^-GFP, MYB36^AAA^-GFP, MYB36^DDD^-GFP with or without EST induction. CASP1-GFP overviews from different plant lines at mature CS stage (16-17^th^ cell) are shown in upper panels. Nuclei signals of MYB36-GFP from the same lines are taken around the onset of CASP1-GFP (7-8^th^ cells) and shown in corresponding panels below. GFP signals are shown in gradient intensity. Scale bars = 20µm

**Movie S1. CASP1-GFP phenotypes of *2-in-Endo* with CIF2 or flg22 treatments. Related to Figure 1**. Total imaging time 5.5 h, each frame taken at 30 min intervals. Scale bars=25µm.

**Movie S2. SGN preferential reporter pPRK1::NLS-3xmVenus in *2-in-Endo.* Related to Figure 2**, Table S1. Total imaging time 8 h, each frame taken at 1 h intervals. Scale bars=25µm.

**Movie S3. FLS2 preferential reporter pWRKY30::NLS-3xmVenus in *2-in-Endo.* Related to Figure 2**, Table S1. Total imaging time 8 h, each frame taken at 1 h intervals. Scale bars=25µm.

**Table S1.**
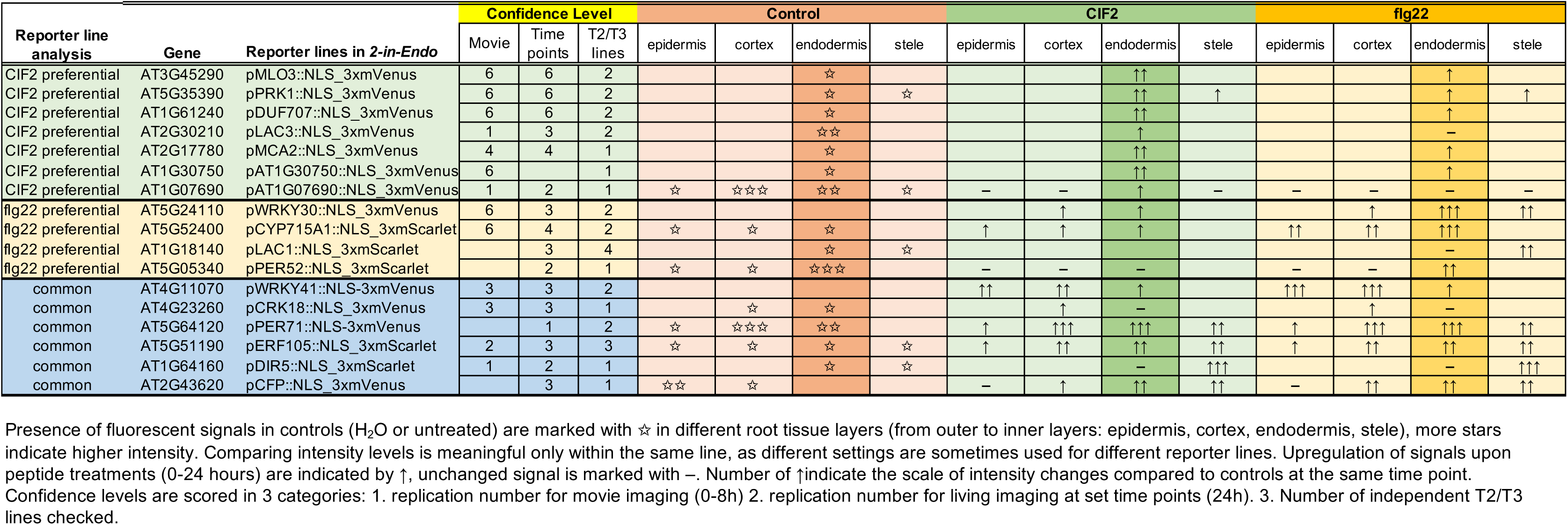
Summary of reporter line analyses. Related to Figure 2.

**Table S2.** T1 phenotype distributions of *myb36 CASP1-GFP* transformed with MYB36 variants (WT, AAA, DDD) either untagged, or with C-terminal mCherry or Turbo-GFP tags. Related to Figure 3.

|  | myb36 CASP1-GFP |  |  |  |  |  |  |  |  |  |
| --- | --- | --- | --- | --- | --- | --- | --- | --- | --- | --- |
| CAPS1-GFP phenotype | untransformed | untagged MYB36 |  |  | mCherry tagged MYB36 |  |  | Turbo-GFP tagged MYB36 |  |  |
|  |  | WT | AAA | DDD | WT | AAA | DDD | WT | AAA | DDD |
| Fully fused "wild-type" |  | 17/20<br>= 85% | 5/20<br>= 25% | 18/20<br>~ 90% | 15/18<br>~ 83% | 6/10<br>= 60% | 13/15<br>~ 86% | 10/10<br>= 100% | 12/14<br>~ 86% | 9/10<br>= 90% |
| Discontinuous "bubbles"and/or patchy |  | 1/20<br>= 5% | 12/20<br>= 60% | 1/20<br>= 5% | 3/18<br>~11% | 3/10<br>= 30% | 2/15<br>~ 13% |  | 2/14<br>~14% | 1/10<br>=10% |
| Weak cytoplasmic or no signal | 20/20<br>= 100% | 2/20<br>= 10% | 3/20<br>= 15% | 1/20<br>= 5% |  | 1/10<br>= 10% |  |  |  |  |

Number of observations matching each phenotype category and percentage in total T1 observations are shown.

**Table S3.**
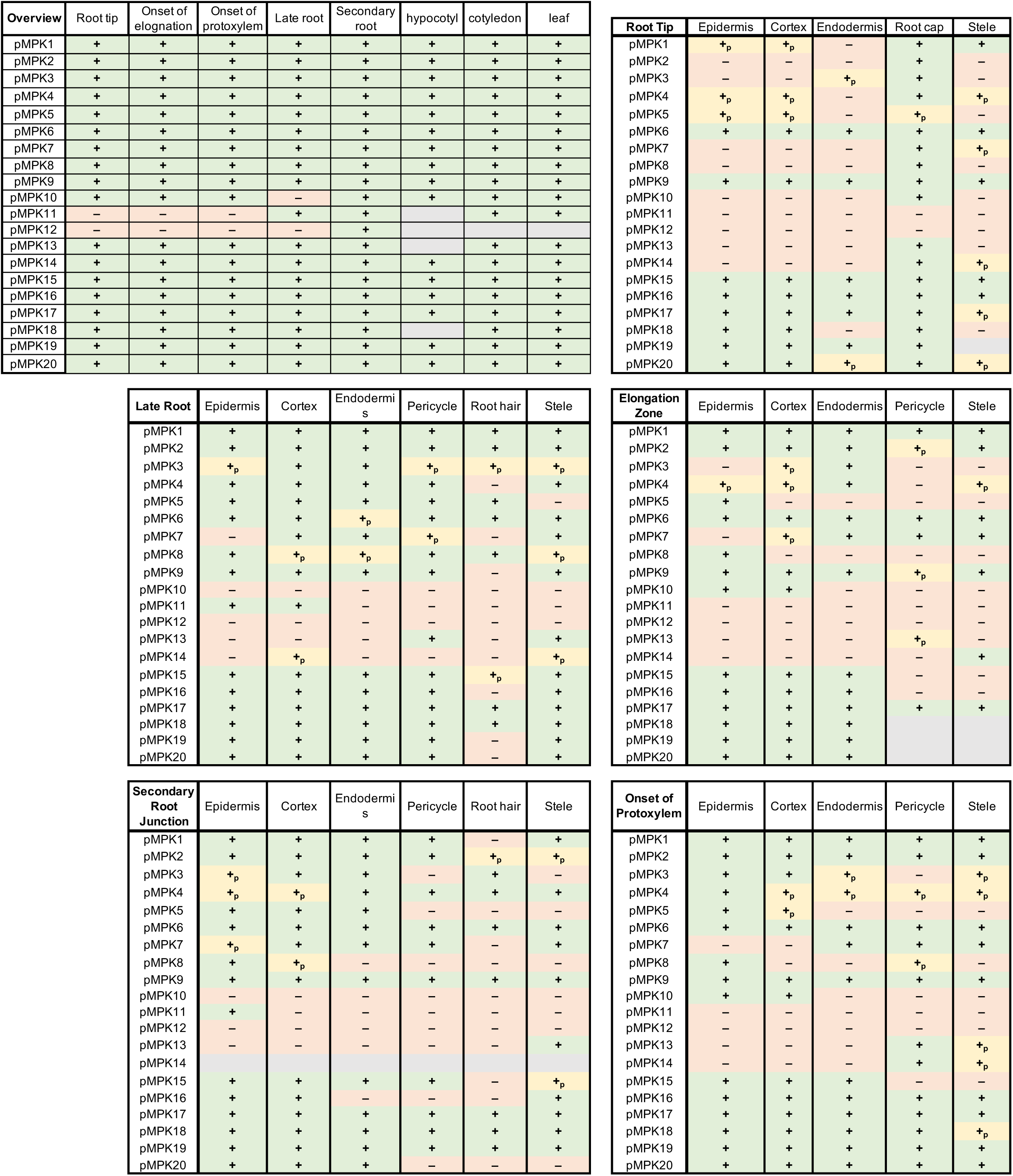
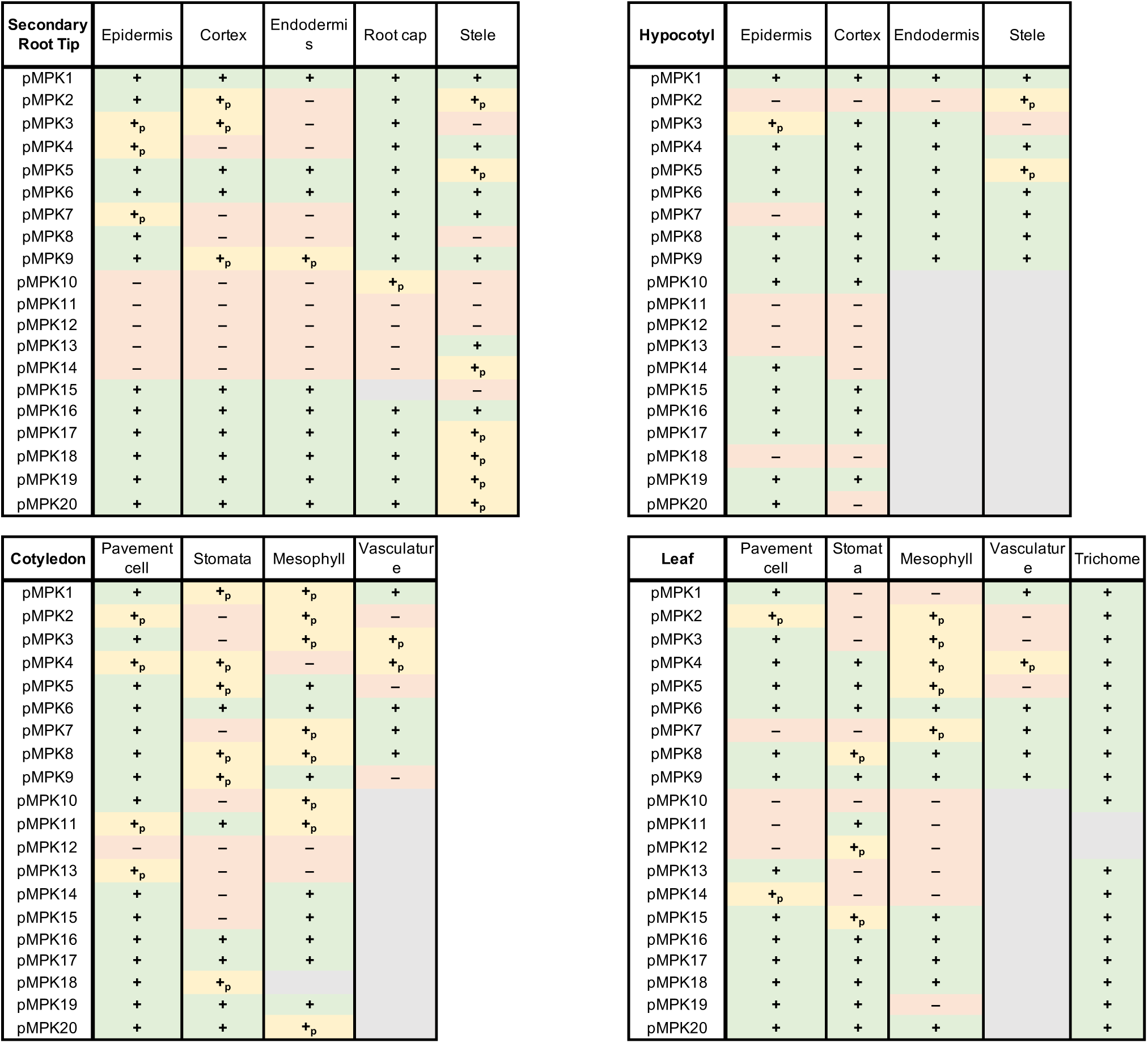
Expression atlas of 20 Arabidopsis MPK in various tissues. Related to Figure 4. Data based on analysing fluorescence signals from individual pMPK::NLS-3xmVenus line (1 to 20) in wild-type (10-day-old roots). Presence (green +), absence (red –) or not observable (grey) of signals are indicated. Not observable cases were due to tissue opacity in live imaging. Overview summarizes expression patterns in multiple plant locations: Root Tip, Elongation Zone (in root), Onset of Protoxylem (7-8th cell after onset of elongation), Late Root (after onset of metaxylem), Secondary Root, Hypocotyl, Cotyledon and Leaf. At each location, MPK expressions were analysed in each cell/tissue types individually. Expression in all cells (green +), expression in some but not all cells (yellow +p), no expression in all cells (red –), not observable (grey).

## STAR Methods

### KEY RESOURCES TABLE

### RESOURCE AVAILABILITY

- **Lead Contact** Further information and requests for resources and reagents should be directed to and will be fulfilled by the Lead Contact, Niko Geldner.
- **Materials Availability** Plasmids and transgenic plant seeds generated in this study will be made available on request, but we may require a payment and/or a completed Materials Transfer Agreement if there is potential for commercial application.
- **Data and Code Availability.** The full RNA-seq dataset will be deposited in GEO. This study does not generate new code. R scripts used to generate graphs or Fiji Macros used for image processing and quantifications will be made available on request.

### EXPERIMENTAL MODEL AND SUBJECT DETAILS

#### Plant material

*Arabidopsis thaliana* ecotype Columbia (Col-0) was used as wild-type controls and the background ecotype for all experiments. The mutants *fls2* (SALK_062054C)^5^, *sgn3-3* (SALK_043282)^6^, *cif1 cif2*^1^ *(Generated by CRISPR Cas9)* and pSGN3::SGN3-mVenus in wild-type^6^ were previously described. *2-in-Endo* line were generated in two steps: 1. pSGN3::FLS2-3myc-GFP was introduced into *fls2* (SALK_062054C) background 2. *cif1 cif2 null* alleles were obtained by CRISPR-Cas9 using a single sgRNA (5’ - gctttggtttaggactggag -3’) that targets both genes.^1, 7^ The *pCASP1::FLS2-3myc-GFP fls2* (SAIL691_C04) described previously^8^ *was crossed to sgn3-3* (SALK_043282) to obtain *pCASP1::FLS2-GFP fls2 sgn3*.

T-DNA mutant seeds mpk2 (SALK_047422C), mpk3 (SALK_151594), mpk6 (SALK_073907), mpk9 (SALK_064439C), mpk16 (SALK_059737C), mpk17 (SALK_020801C), mpk19 (SALK_075D213C), mkk4 (SALK_0188040C), mkk5 (SALK_047797C) and mkk9 (SAIL_60_H06) were provided by the Nottingham Arabidopsis Stock Centre (NASC). Homozygous double mutants mpk2 mpk16 and mkk4 mkk5 were generated by crossing the above indicated T-DNA mutants. mkk1 (Salk_027645) mkk2 (Sail 511_H01) and mkk3 (SALK_051970)^9^ were obtained from Jean Colcombet, mpk3 mpk6 pMPK6::MPK6^YG^ and mpk3 mpk6 pMPK3::MPK3^TG^ were obtained from Zhang Shuqun.^10, 11^

The published constructs pCASP1::CASP1-GFP with FastRed selection cassette^12^ and pGPAT5::mCitrine-SYP122^13^ were introduced into respective backgrounds by floral dipping. For pathway specific reporters, promoter fusion constructs in Table S2 were introduced into the *2-in-Endo* background. For MPK expression atlas (Table S3), promoter fusion constructs pMPK1-20::NLS-3xmVenus were introduced into wild-type background.

#### Plant growth conditions

Plant seeds were surface sterilized in 70% ethanol + 0.05% Tween20 for 7-10 min, then washed in 96% ethanol and dried in sterile conditions. Seeds were stratified for 2 days at 4 °C in the dark on half-strength Murashige and Skoog (½MS) + 0.8% Agar (Roth) plates containing 500 mg/L MES (Duchefa). Seedlings were grown vertically for 5-6 days at 23°C under continuous light before analysis.

### METHOD DETAILS

#### Plasmid construction and Plant transformation

The In-Fusion Advantage PCR Cloning Kit (Clontech) and Multisite Gateway Cloning Technology (Invitrogen) were used for generation of constructs.

All inducible constructs (ind) were assembled by triple Gateway recombination reaction (LR) between Entry clones P4-pELTP (LTPG15)-XVE-P1r (promoter)^14^, L1-HopAI1-L2, L1-MPKa-L2 or L1-MKKa-L2 (Gene-of-interest), R2-3xFlag-L3 (C-terminal tag) into the destination vector pF7m34GW,0 containing a FastRed plant selection cassette with a 35S terminator. cDNA entry clones of HopAI1 and HopAI1^H103A^ were synthesized by Genscript^15^; MPK6a (D218G E222A)^16^ was obtained from Jean Colcombet; MPK2a (Q188G E192A), MKK1a (T218E S224D), MKK2a (T229D T235E), MKK3a (S245E T241D), MKK4a (T224D S230E), MKK5a (T215E S221E), MKK6a (S221D T227E), MKK7a (S193E S199D), MKK8a (S195D S201E), MKK9a (S195E S201E), MKK10a (S197E) were cloned from Col-0 cDNA and mutations obtained through site-directed mutagenesis.

All MYB36 constructs were assembled by triple Gateway recombination reaction between Entry clones P4-pMYB36(4kb)-R1 (native promoter), L1-MYB36^WT^-L2, L1-MYB36^AAA^-L2 or L1-MYB36^DDD^-L2 (Gene-of-interest), and R2-MYB36-native terminator(250bp)-L3, R2-mCherry-4G-L3 or R2-SuperfolderGFP-TurboID(BirA*)-6His-TEV-3FLAG-L3 (Terminator or C-terminal tag) into destination vectors pF7m34GW,0 containing a FastRed plant selection cassette or pFG34GW, containing a FastGreen plant selection cassette. Genomic DNA entry clones of wildtype MYB36 were cloned from Col-0 gDNA, and mutants MYB36^AAA^ (S18A, S146A, S169A) and MYB36^DDD^ (S18D, S146D, S169D) were synthesized by Genscript.

The pathway specific reporter constructs (Table S1) and MPK reporter constructs (Table S3) were made firstly by generating promoter entry clones using In-Fusion: the endogenous promoter sequences (around 2kb before start codon) were cloned from Col-0 gDNA or JAtY clones. The reporter constructs were then assembled by double Gateway recombination reaction between a promoter entry clone and L1-Nuclear Localisation Signal (NLS) fused to a triple mVenus fluorescent tag (3xmVenus) -L2 or L1-NLS-3xmScarlet-L2 into the destination vector pFG24GW (FastGreen), pFR7m24GW (FastRed) or pB7m24GW (BASTA selection). See Key Resources Table for details.

All plasmids were transformed by heat shock into *Agrobacterium tumefaciens* GV3101 strain with pSoup plasmid (pMP90) and then transformed into the corresponding plant lines by floral dip method.^17, 18^ Based on the plant selection marker, around 10-30 transgenic lines were selected and characterized at T1 generation. From these, 2-6 independent transgenic lines (T2) were isolated for further characterization. Independent homozygous lines at T3 were confirmed by non-segregation of the selection marker and were used for western blot experiments or as background for further introduction of constructs.

#### Seedling growth and treatments

The flg22 peptide from *Pseudomonas aeruginosa* (QRLSTGSRINSAKDDAAGLQIA) was ordered from EZBioLab; and CIF2 peptide (DY*GHSSPKPKLVRPPFKLIPN, *: sulfated tyrosine) from Peptide Specialty Laboratories GmbH (https://www.peptid.de/). Peptides were dissolved in deionized MilliQ sterile water at the stock concentration of 1mM and diluted in melted ½ MS agar medium to indicated concentrations. Peptide treatments: Seedlings were grown vertically for 4-5 days, then transferred with care onto plates containing peptides or H_2_O control to grow vertically for 1 day before analysis.

For time lapse imaging and movies with peptide treatments, 3-day-old seedlings grown vertically on ½ MS plates were transferred to hydroponic system (a 12-well Multiwell plate (CytoOne) with 6.5 mL liquid ½ MS in each well) to grow for 2 days. These 5-day-old seedlings were then transferred inside a chambered coverglass (Thermo Scientific) and then covered with a ½ MS agar block containing 1 µM peptide. 50 μL of liquid ½ MS containing the same peptide concentration is added through a channel in the middle of agar block, directly reaching the roots.

10 mM stock of EST (Sigma, solvent DMSO) and NM-PP1 (Adipogen, solvent DMSO) was diluted to 5 µM in melted ½ MS agar medium, and 0.05% v/v DMSO was used as control treatments. For estradiol (EST) treatments, seedlings were grown vertically on EST or DMSO plates for 5 days or transferred to EST or DMSO to grow for 1 day. For NM-PP1 treatments, seedlings were grown vertically on ½ MS plates for 4 days and transferred to NM-PP1 or DMSO plates to grow vertically for 1 day.

#### Fluorescence Microscopy

Imaging was performed using Leica SP8, Leica Stellaris, or Leica Stellaris 5 WLL IR confocal laser scanning microscopes and Leica Thunder light Microscope (DM6BZ). Images were taken with a 63 × water immersion objective (Leica SP8, Leica Stellaris) at zoom factor 1 for overviews and zoom factor 5 for surface views. Time lapse imaging of Figure 2F and Movies 2,3 were taken with a 63 × oil immersion objective (Leica Stellaris 5 WLL IR). Time lapse imaging of Figure S3D and Movie 1 were taken with a 40 × oil immersion objective (Leica Stellaris). Root scans for Fluorol Yellow (suberin staining) and pGPAT5:mCitrine-SYP122 (suberin synthesis) were acquired by Leica Thunder light Microscope with 10 × dry objective and tile-scan (10% overlap). Unless indicated otherwise, confocal images comparing genotypes/treatments were taken following the “four identical criteria”: same position in the roots, the same laser detection intensity, the same laser scanning area (zoom), and the same interval of slices for Z stack projection. For overview images, maximum projection of z-stacks (step size = 1 µm) are shown. For surface view images, maximum projection of z-stacks (step size = 0.36 µm) are shown.

The excitation and detection windows were set as following. For live imaging, GFP (488 nm, 495-550 nm), PI (488 nm, 600-690 nm), mVenus (514nm, 520-565nm), mCitrine (514 nm, 555-570 nm), mScarlet (561 nm, 580-640 nm), mCherry (561 nm, 585-680 nm). For time lapse imaging with Leica Stellaris 5 WLL IR, mVenus (507 nm, 520-560 nm), mScarlet (555 nm, 580-600 nm). For fixed samples, sequential scanning was used to avoid interference between the following fluorescence channels: calcofluor white (405 nm, 415-435 nm), GFP (488 nm, 495-525 nm), basic fuchsin (561 nm, 600-650 nm). For suberin analysis on Thunder, Fluorol yellow (488nm, 500-550nm), mCitrine (488nm, 500-550nm).

#### Propidium iodide (PI) penetration assay

PI penetration assay assess the stage of the formation of a functional CS diffusion barrier.^19, 20^ Seedlings were incubated in 10 µg/ml PI for 10 mins and rinse in H_2_O before imaging. PI block is identified as the endodermal cell achieves the exclusion of PI signal from the inner side. The number of endodermal cells is counted from the onset of elongation until PI block. For roots with no functional barrier, PI is not blocked near the root-hypocotyl junction, and thus are shown as individual data points in the category “near hypocotyl” and are excluded from numerical statistical test.

#### Suberin quantification assay

Suberin was stained with Methanol-based Fluorol Yellow protocol.^1, 8^ Seedlings were fixed and cleared in methanol for at least 3 days at 4 °C, exchanging the methanol at least once. The cleared seedlings were transferred to a freshly prepared solution of Fluorol Yellow 088 (0.01% in methanol) and incubated for 1 hour at room temperature or overnight at 4 °C in the dark. The stained seedlings were rinsed in methanol and transferred to a freshly prepared solution of aniline blue (0.5% in methanol) for 1 hour counterstaining in the dark. Finally, the seedlings were washed briefly in water before mounting and imaging with the Leica Thunder light microscope.

#### Fixation and lignin staining

Fixation and staining were performed with adapted Clearsee protocol.^21, 22^ Briefly, seedlings were fixed in 4% paraformaldehyde PBS solution at 4 °C overnight, using 6-well or 12-well plates (CytoOne), then washed twice for 1min with PBS. Once fixed, seedlings were cleared in Clearsee solution for at least 24h under gentle shaking. Fixed and cleared samples were then incubated overnight in a Clearsee solution supplemented with 0.2% Basic Fuchsin and 0.1% Calcofluor White for combined cell wall and lignin staining. Once the dye solution removed, samples were rinsed overnight in Clearsee twice before mounting and observation with confocal microscopes.

#### Electron microscopy for ROS and lignin detection

Visualization of H_2_O_2_ (ROS) around the CS was performed with the cerium chloride assay described previously.^1, 7^ This detection method is based on Ce^3+^ ions reacting with H_2_O_2_, forming electron-dense cerium perhydroxide precipitates, which are detected by transmission electron microscopy. Briefly, seedlings were incubated in 50 mM MOPS buffer (pH 7.2) containing freshly prepared 10 mM cerium chloride (CeCl_3_) (Sigma) for 45 min. Seedlings were then washed twice in MOPS buffer for 5 min and fixed in glutaraldehyde solution (EMS) 2.5% in 100 mM phosphate buffer (pH 7.4) for 1 h at room temperature. Then, they were postfixed in osmium tetroxide 1% (EMS) with 1.5% potassium ferrocyanide (Sigma) in phosphate buffer for 1 h at room temperature. The plants were rinsed twice in distilled water and then dehydrated in ethanol solution (Sigma) at gradient concentrations (30%, 40 min; 50%, 40 min; 70%, 40 min; 100%, 1 h x 2). This was followed by infiltration in Spurr resin (EMS) at gradient concentrations (Spurr 33% in ethanol, 4 h; Spurr 66% in ethanol, 4 h; Spurr 100%, 8 h, twice) and finally polymerized for 48 h at 60 °C in an oven. Ultrathin sections (50 nm) were cut transversally at 1.5, 2 or 3mm from the root tip on a Leica Ultracut (Leica Mikrosysteme GmbH) and picked up on a nickel slot grid 2 × 1 mm (EMS) coated with a polystyrene film (Sigma).

Visualization of lignin deposition around the CS and cell corner was done using permanganate potassium (KMnO_4_, Sigma) staining protocol.^23, 24^ The sections were post-stained with 1% of KMnO_4_ in H_2_O for 45 min and rinsed several times with H_2_O.

Micrographs were taken with a transmission electron microscope FEI CM100 (FEI) at an 80 kV acceleration voltage with a TVIPS TemCamF416 digital camera (TVIPS GmbH) using the software EM-MENU 4.0 (TVIPS GmbH). Panoramic were aligned with the software IMOD.^25^

#### RNAseq experiments and analysis

Sample preparation for RNAseq experiments was as previously described.^1^ Briefly, wild-type, *sgn3*, *fls2* and p*CASP1::FLS2-GFP fls2* seeds were grown vertically on a sterile mesh on solid ½ MS plates for 5 days and transferred onto fresh ½ MS solid medium containing 1 μM peptides and mock plates (mock and peptide treatments always transferred in parallel). After 30-, 120- and 480-mins incubation on plates, whole roots from 2 square plates (50 ml) were cut and collected for each sample, then immediately frozen in liquid nitrogen. Three biological replicates for each sample were conducted on three different days. Total RNA was extracted with a TRIzol-adapted ReliaPrep RNA extraction kit (Promega).

Libraries were prepared by Lausanne Genome Technology Facility following the same protocol of Fujita *et al*.^1^ RNA quality control was performed on a Fragment Analyzer (Advanced Analytical Technologies). 1000 ng of total RNA was used to prepare RNA-seq libraries with the Illumina TruSeq Stranded mRNA reagents (Illumina) on a Sciclone liquid handling robot (PerkinElmer) using a PerkinElmer-developed automated script. The resulting library was used for cluster generation with the Illumina TruSeq SR Cluster Kit v4 reagents and sequenced on the Illumina HiSeq 2500 using TruSeq SBS Kit v4 reagents. The Illumina Pipeline Software version 2.20 was used to process sequencing data.

Lausanne Genomic Technologies Facility performed the data processing using their in-house RNA-seq pipeline, as described in Fujita *et al.*^1^ Briefly, purity-filtered read trimming for adapters and low-quality sequences was done with Cutadapt (v. 1.8)^26^ and removal of reads matching ribosomal RNA sequences with fastq_screen (v. 0.11.1). Low complexity reads were filtered with reaper (v. 15-065)^27^ . Cleaned reads were aligned against *A.thaliana* TAIR10 genome using STAR (v. 2.5.3a)^28^ and read counts per gene locus were obtained with htseqcount (v. 0.9.1)^29^ using *A. thaliana* TAIR10 Ensembl 39 gene annotation. RSeQC (v. 2.3.7)^30^ was used to evaluate the quality of the data alignment.

Statistical analysis was performed for genes in R (3.5.3). Genes with low counts were filtered out according to the rule of one count per million (cpm) in at least one sample. Library sizes were scaled using TMM normalization and log-transformed into counts per million or CPM (EdgeR package version 3.24.3).^31^ Data was corrected for the experimental batch effect using removeBatchEffect function (limma). Statistical quality controls were performed through pairwise sample correlations, clustering and sample PCA using batch corrected normalized data. Differential expression was computed with limma-trend approach by fitting all samples into one linear model.^32^ The experimental batch factor was added to the model. Moderated t-test was used for each pairwise comparison (peptide-treated vs H_2_O-treated) per time point. The adjusted p-value is computed by the Benjamini-Hochberg method, controlling for false discovery rate. To determine genes significantly regulated by CIF2 or flg22 (* Figure 2, S2), the adjusted p-value was below 0.05 (P<0.05).

Heatmaps gene lists in Figure 2C, D were selected based on published genes involved in lignin and suberin biosynthesis.^1, 33^ Figure 2 E genes were based on the intersection of datasets by Liberman *et al* (RNAseq)^3^ and Kamiya *et al* (microarray)^34^ that report significant downregulation in *myb36* mutants compared to wild-type. *myb36* vs wild-type regulation heatmap (left) was generated using Liberman *et al*’s data.^3^ Figure 2B genes were based on genes significantly upregulated by flg22 in whole seedlings.^15^ WT treated with flg22 heatmap (left) was generated using li *et al*’s data. ^15^

GO-enrichment analyses were performed by clusterProfiler R package (3.18.1) with “Biological process”, adjusted p-value use Benjamini-Hochberg method, cutoff 0.05. GO annotations use org.At.tair.db R package (3.12.0). Gene ratio represents the gene counts in a GO category that are differentially regulated by a pathway divided by the total number of genes in that GO category.

#### Western Blots

For Figure 4A, seedlings were grown vertically for 5 days on sterilized mesh on solid ½ MS media. The seedlings were treated by adding 20 ml liquid ½ MS containing respective peptides or H_2_O control to submerge the roots. At 5, 10, 15min liquid was drained and roots were cut and collected. For Figure S5 F, G, seedlings were grown for 5-6 days on sterilized mesh on solid ½ MS media supplemented with EST or DMSO.

Root tissues collected were immediately frozen in liquid nitrogen and grounded by Tissue Lyser II (Qiagen). Extraction buffer (50 mM Tris HCl pH7.5, 150 mM NaCl, 0.5 mM EDTA, 1% Triton x-100, 50 mM ϕ3 glycerophosphate, 1 mM PMSF, 100 uM Na orthovanadate, 5 mM Na fluoride, 2.5 mM MgCl2, 1x cOmplete ™ EDTA-free Protease Inhibitor Cocktail) was added to the frozen samples. The samples were briefly mixed by vortex and centrifuged at 15000 rpm for 15 min at 4°C. The supernatant was transferred in new tubes and measured the concentration by Bradford method (Thermo Fisher Scientific). After adjusting concentration, the proteins were denatured by addition of 3x (v/w) 1x NuPAGE LDS sample buffer + 10 mM DTT (Invitrogen, NP0007) and heated for 5min at 95°C.

Equal amount of protein was loaded (∼20-40 ug/lane) and separated on 10% or gradient (4-12%) pre-cast acrylamide gels (Eurogentec). After electrophoresis, the separated proteins were transferred onto PVDF membrane (Immobilon-E, Merck-Millipore) using Pierce FastBlotter G2 semi-dry blotting. (ThermoFisher).The blot was blocked in 1% blocking buffer in TBS (western blocking reagent, Roche) for 1 h at room temperature before incubation with primary antibody: anti-p44/42 (1:1000, Cell Signaling Technologies #4370), anti-MPK6 (1:10000, Sigma), anti-FLAG conjugated M2 Peroxidase (1:500, Sigma), anti-Actin (1:2000, Sigma) in 0.5% blocking buffer + TBS-0.05% Tween overnight at 4°C. After washing the blot 2-3 times with 1x TBS + 0.1% Tween (TBS-T), the blot was treated with secondary antibody (anti-rabbit, 1:10000, Agrisera for anti-p44/42; anti-rabbit HRP ECL 1:30000 for anti-MPK6, anti-mouse HRP, 1:5000, Promega for Actin) in 0.5% blocking buffer + TBS-0.05% Tween for 2 h and then washed 5-7 times with TBS-T. For loading control detection of MPK6 and Actin, the original membrane was stripped (Thermo Scientific). Signal was detected with SuperSignal West Pico Plus and Femto kits (Thermo Scientific).

### QUANTIFICATION & STATISTICAL ANALYSIS

All fluorescence microscopy images were processed using Fiji (ImageJ2 Version:2.9.0/1.53t) with Fiji Macros to semi-automate the processing and quantification.^35^ Macros will be made available on request.

Percentage (%) of gaps in CASP1-GFP (Figure 1E, S1H) are based on maximum projections of surface view images (GFP channel) after thresholding to distinguish background from signal. Quantification represents the percentage of uncovered area (background) of a fused CASP1-GFP belt reconstructed from connecting individual domains using a selection Bush tool. Fully connected CASP1-GFP yield 0% of gaps.

Number of holes per 100 µm of CASP1-GFP (Figure S1C, 3E, 4E, 5B, S5C) are based on maximum projections of overviews (GFP channel) after thresholding to distinguish background from signal. Steps for quantification are detailed in Figure S1B. Each data point represents the average number of holes per 100 µm calculated from a single overview image.

Quantifications of total grey intensity of nuclei signal (Figure 2F) are calculated from Sum Slice Projection of overview images at each hour. This method sums the intensity values at each pixel location throughout the stack and projects the sum onto a 2D image. This is useful for visualizing the total intensity contribution from all slices in the stack, especially in the cases of overlapping nuclei. Thresholding and ROI selection were then applied to isolate nuclei signals from noise and background. Similar procedures were used for Figure 6 to calculate total grey intensity of MYB36 nuclei signal from Maximum projection of overview images.

For quantitation of CASP1 domain properties: Number of fragments per 100 µm (Figure 3C) and the Average size of fragments (Figure S3C) were based on the same data points from maximum projections of surface view images. Thresholding followed by Particle Analysis (no minimum size) were used to quantify the number of fragments and the average size of fragments in a single image. The length of CASP1 in µm for a single image were measured and used to calculate Number of fragments per 100 µm.

Quantification of CS coverage (Figure 5D, S5D, 6B) is represented by the Percentage of CASP1 length coverage normalized to wild-type. Total CASP1 length (measured after applying skeletonize (Fiji)) of each overview image (maximum projection) were measured and then normalized to the average CASP1-GFP length in wild-type roots from the same experiments to calculate the percentage of CS coverage for each data point.

All graphs and statistical analyses were performed using R 4.0.4 (http://www.R-project.org/). For multiple comparisons, ANOVA followed by Tukey’s Honestly Significant difference (HSD) test were applied when linear model assumptions were met. Unbalanced Tukey test were used for unequal replication. For comparing the mean of one sample with a control, Student’s t-test were used equal replicates, Welch t-test for unequal replicates. For example, the analysis of suberization comparing DMSO with EST treatments, t-tests were performed for each zone separately, and letters indicates significant differences for a given zone (a, b or a’, b’). In box plots, boxes span the first to third quartiles, with the bold line representing the median. Whiskers indicate maximum and minimum values, except outliers. Bold line in dot plot represents the mean. Each point in the box plot or dot plot represents a data point from an individual root (n). Statistical parameters, error bars and n are reported in the figure legends.

